# Reversed functional gradient in primate prefrontal cortex: posterior dominance and frontopolar deactivation

**DOI:** 10.1101/2025.06.16.659673

**Authors:** Kei Watanabe, Masayuki Hirata, Takafumi Suzuki

## Abstract

The frontopolar cortex (FPC) is thought to coordinate the more posterior lateral prefrontal cortex (LPFC) during complex, non-routine behaviors through high-level functions such as management of multiple goals, exploration, and self-generated decision-making. However, direct neurophysiological comparisons with other prefrontal regions are lacking, leaving the FPC’s putative dominance untested. Contrary to this view, our comparison of neuronal activity across the full anteroposterior LPFC in macaques during six distinct tasks probing these functions revealed a posterior-to-mid LPFC dominance, with resource-allocation, novelty-detection (including reward prediction error), and modality invariant decision-monitoring signals all showing a common posterior bias. In contrast, regardless of task demands, the FPC’s strongest encoding was about the most recently executed action, and it displayed minimal object selectivity, even when objects were task-critical. We identified a turning point in this graded posterior-to-anterior transition from task-positive to task-negative regions around the border between the anterior and middle thirds of the LPFC. These findings challenge the prevailing notion that the LPFC is anterior-dominant across primate species, and provide evolutionary constraints on theories of human prefrontal organization.

## INTRODUCTION

While an understanding of the functional organization of the lateral prefrontal cortex (LPFC) is key to unraveling the neural basis of intelligent behavior, consensus remains elusive. A prominent view suggests an anterior-dominant functional gradient in the LPFC in which more anterior regions, including the frontopolar cortex (FPC; Brodmann area 10), play progressively abstract, integrative, and supervisory roles over posterior regions^1–3^. Others argue that the apex of the prefrontal hierarchy lies in the mid-dorsolateral PFC^4,5^ or that the LPFC does not follow a fixed hierarchical structure^6^. Thus, it is critical to understand frontopolar function to elucidate how the broader LPFC orchestrates complex cognition, yet this understanding remains poorly developed. In macaques, a key model for human brain function, FPC studies are scarce, and the results of lesion^7–10^ and neural recording studies^11–14^ are not straightforward. Focal FPC lesions do not cause major deficits in standard cognitive tasks involving memory, value, rules, or strategies^7–9^, and FPC neurons encode these parameters weakly^11,12^. Instead, the monkey FPC appears to be selectively engaged in more complex, non-routine situations, including (i) managing multiple goals by allocating cognitive resources (cognitive orchestration)^8^, (ii) exploring novel environments (rapid novel learning)^7,13^, and (iii) monitoring ‘self-generated decisions’ to integrate past experiences with current cues for guiding future behavior^11^. These capacities are considered to be key facets of an overarching function: adaptive control of behavior in dynamic, integrative contexts^15^.

However, this adaptive role is not unique to the FPC. The mid-to-posterior LPFC, along with medial and orbital PFC and subcortical structures, is also engaged in cognitive orchestration^16^, novel learning^17–20^, and decision monitoring^21–23^. Because systematic neurobiological comparisons with other prefrontal regions are lacking, the contribution of the FPC may have been overemphasized, leaving its putative dominance untested. Moreover, FPC studies in monkeys are scarce, employ a limited range of experimental conditions, and often analyze only neurons pre-screened for robust task-related responsiveness^11,13^, making it premature to conclude that the FPC is genuinely critical for these processes. For instance, evidence for the monkey FPC’s role in cognitive orchestration comes from a single lesion study (where the effect size is compared only to that of the posterior cingulate cortex)^8^, and underlying neuronal mechanisms are unexplored. In self-generated decision monitoring, the only prior FPC recording study focused exclusively on spatial tasks, overlooking object-based decision-making^11^. These limitations warrant a systematic, full-sample comparison of neural activity across the entire anteroposterior LPFC to determine how these functions are distributed.

In this study, we examined neuronal activity across the LPFC while monkeys performed six tasks designed to probe the proposed functions of the FPC. These comprised (i) unimodal and multimodal dual tasks taxing cognitive orchestration (Experiments 1–2), (ii) two types of rapid novel learning tasks engaging exploratory behavior (Experiments 3–4), and (iii) spatial and object self-generated decision tasks requiring integration of internal and external information for behavioral adaptation (Experiment 5). All tasks, except for the spatial self-generated decision task, are new to FPC electrophysiology. Since they use both spatial and object-based information and multiple motor outputs (saccadic and manual responses), these experiments provide a rigorous test of whether the FPC plays a dominant role in the broader LPFC. We found that adaptive control signals for these key functions peak in the posterior-to-mid LPFC, with resource-allocation, novelty-detection (including reward-prediction error), and modality invariant decision-monitoring signals all showing a common posterior bias. By contrast, FPC activity was markedly inflexible: it primarily encoded the spatial location of just-executed actions and exhibited only minimal object selectivity—even during object-based novel learning and self-generated decision monitoring, where object information was critical.

## RESULTS

### Localization of neural activity

In each daily session, we inserted a linear multi-electrode probe (U-probe) into each of the two recording chambers: one targeting the FPC and the other the mid-to-posterior LPFC (**Figure 1A**). We used a grid system (**Figure 1B**) for precise probe placement. To reconstruct recording locations, we acquired structural MRI images with the grid filled with a contrast agent. We then determined the stereotaxic coordinates where each probe contacted the cortex, designating these as the recording site for the neurons it recorded. For example, the probe track in **Figure 1B** (red line, top left panel) contacted the cortex (red diamond) at stereotaxic coordinates of anteroposterior (AP) 46.5 mm and mediolateral (ML) 5.0 mm in the left FPC of monkey Um (top right and bottom panels). **Figure 1C** shows all reconstructed recording sites within the FPC and mid-to-posterior LPFC chambers of this monkey, aggregated across experiments. **Figures 1D-1G** show the recording sites for the remaining three monkeys (4 hemispheres). **Figures 1H-1L** show the recording sites organized by experiment. Neurons recorded from the FPC chamber are referred to as FPC neurons, and those from the mid-to-posterior LPFC chamber are pos-PFC neurons. Beyond these labels, we also examined changes in neuronal activity in 1-mm increments along the AP axis. We recorded all neurons encountered without any preselection.

**Figure 1.**
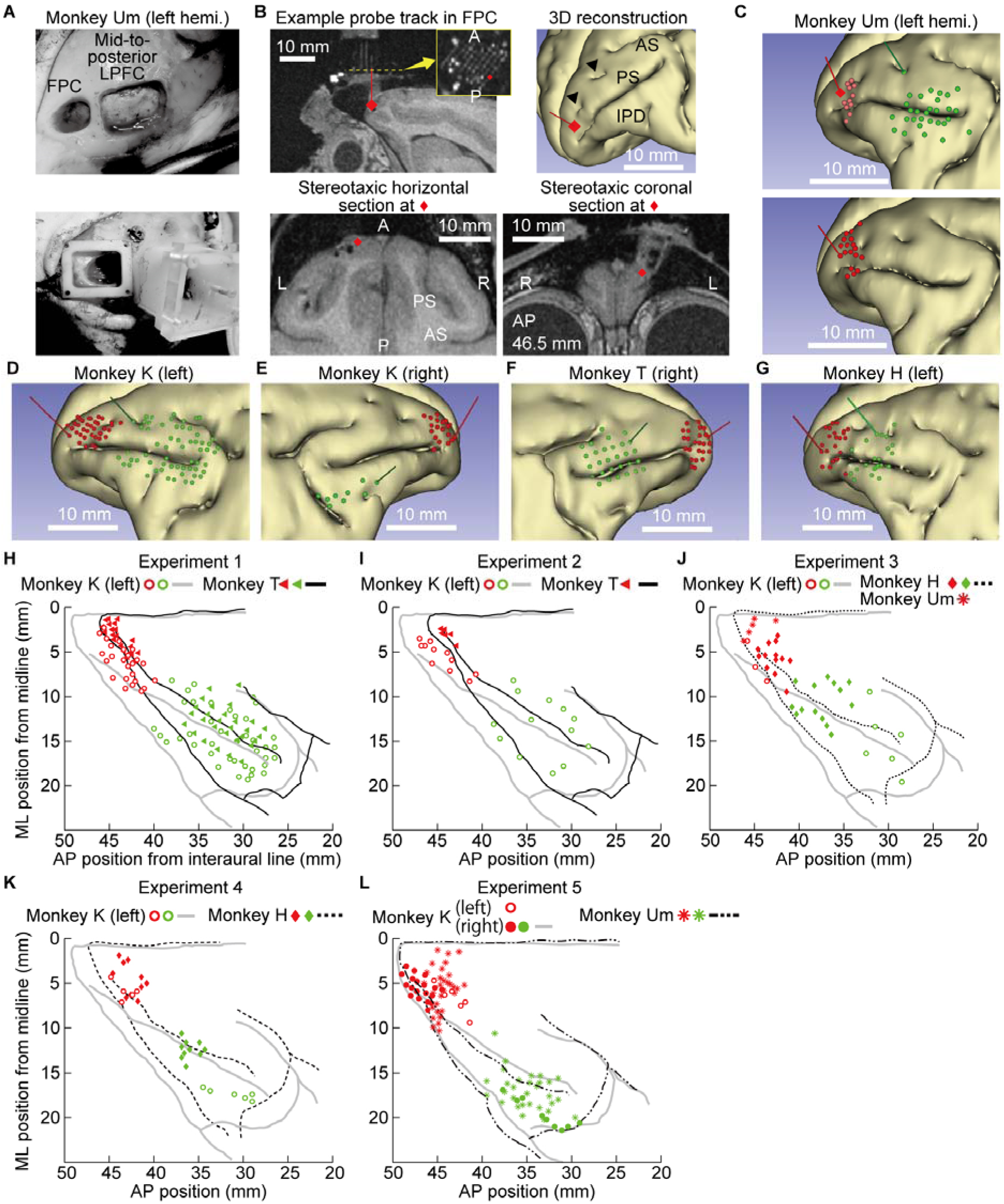
Recording sites. (**A**) Craniotomies (top panel) and recording chambers (bottom) over the FPC and mid-to-posterior LPFC in monkey Um. Hemi., hemisphere. (**B**) Example probe track in the FPC in monkey Um. Top left: A probe inserted through the highlighted grid hole (red) reaches the cortical surface at the red diamond. The white horizontal bar represents 10 mm. Inset: A 1.5× enlarged horizontal section of the grid along the yellow-dotted line. The two larger light spots outside the grid are screw holes for securing it to the chamber. Top right: The same probe trajectory (red line) and cortical entry point (red diamond) shown on a 3D-reconstructed brain of this monkey. PS, principal sulcus; AS, arcuate sulcus; IPD, inferior principal dimple. Of the two black triangles, the rostral triangle indicates the anterior supraprincipal dimple (aspd) and the caudal triangle indicates the posterior supraprincipal dimple (pspd). Bottom panels: Stereotaxic horizontal and coronal sections at the red diamond. A, anterior; P, posterior; L, left; R, right. (**C**) Top: Reconstructed recording sites in monkey Um. The brain is aligned to stereotaxic coordinate space and tilted 30° toward the viewer around the AP axis. Green and red diagonal lines indicate angles of probe insertion in each chamber. Bottom: Because the FPC chamber was relocated once in this monkey, we repeated the reconstruction, based on updated MRI images. Purple and red circles indicate recording sites in the FPC chamber; green circles indicate those in the mid-to-posterior LPFC (pos-PFC) chamber. (**D** to **G**) Same as in **C**, but for the remaining three monkeys, plotted on each monkey’s brain. (**H**) Recording sites for Experiment 1, plotted on a top-down view of the stereotaxically positioned brains of monkeys K (gray outline) and T (black outline). The x-axis and y-axis correspond to the standard AP and ML coordinates, respectively, relative to the interaural midpoint (origin). Right-hemisphere data (monkey T) were flipped to appear as left. Circles and triangles represent sites in monkeys K and T, respectively, with color coding as in **D**. (**I** to **L**) Same as in **H**, but for Experiments 2 – 5. Note that in **j** and **l**, of the three hemisphere outlines, the one with the fewest recording sites was omitted for visibility.

### Experiment 1: Testing the cognitive orchestration hypothesis in the monkey FPC

To examine the FPC’s contribution and to characterize the anteroposterior gradient in cognitive orchestration, we recorded the activities of 2072 single neurons (FPC, *n* = 853; pos-PFC, *n* = 1219; see **Table S1** for the number of neurons recorded in each monkey) in two monkeys performing a dual task (**Figure 2**). The task, as in our previous studies^16,24^, required simultaneous performance of a manual visuospatial attention task and a memory-guided saccade (MGS) task. Recording sites spanned AP 26.5 to 46.1 mm (**Figure 1H**).

**Figure 2.**
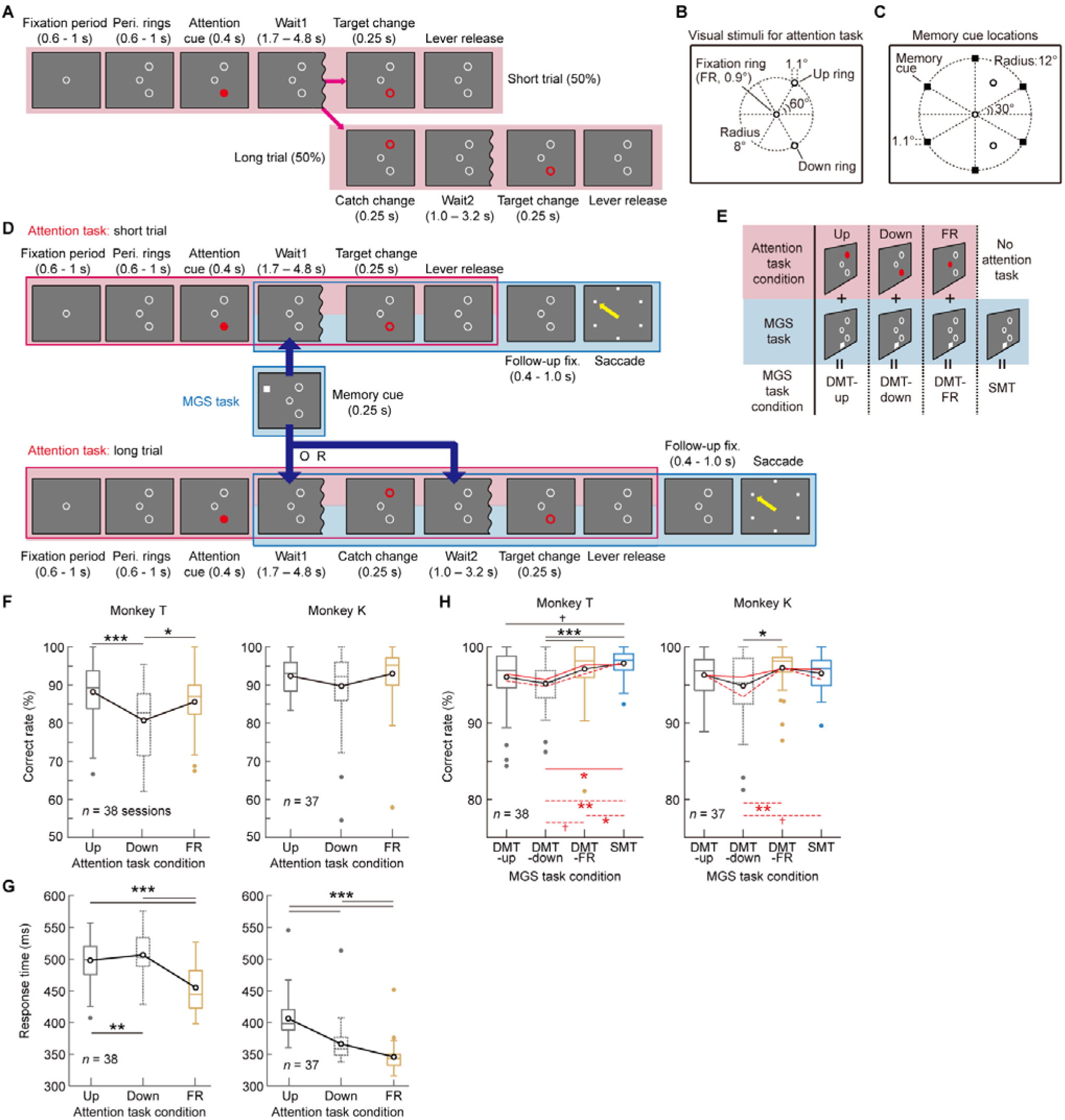
Experiment 1: Behavioral task and performance. (**A**) Event sequence of the attention task. (**B**) Location of visual stimuli in the attention task. (**C**) Location of memory cue presentation. (**D**) Addition of MGS task by inserting a memory cue (middle row) into either the short trial (top row) or long trial (bottom row) of the attention task. (**E**) Combination of the attention and MGS tasks under three DMT conditions and one SMT condition. (**F**) Behavioral performance. Distribution of session-by-session percent correct rates in the three attention task conditions for monkeys T (left) and K (right), shown as boxplots. In each box, the horizontal line and open circle represent the median and mean, respectively. The box edges mark the lower and upper quartiles; whiskers extend to the most extreme values within 1.5 interquartile ranges. Colored dots indicate outliers. Asterisks indicate *p*-values. * *p* < 0.05, ** *p* < 0.01, *** *p* < 0.001 (Holm-Bonferroni corrected). (**G**) Same as in **F**, but for session-by-session mean RTs. (**H**) Distribution of session-by-session percent-correct rates in SMT (cyan) and three DMT conditions for monkeys T (left) and K (right). The solid black line graph shows the mean values when all trials were included (regardless of memory delay length), while the solid and dashed red line graphs show those for trials with short (≤ 3.0 s) and long (> 3.0s) memory delays, respectively (i.e., duration between memory cue offset to go-signal). Lower solid and dashed red horizontal lines indicate the results of statistical tests for short and long memory delay trials, respectively.

Trials of the dual task started with the attention task, initiated by the monkey’s lever-press, which triggered the onset of a central fixation ring (FR). After a brief fixation (**Figure 2A**, ‘fixation period’; 0.6-1.0 s), two peripheral rings (Up and Down rings; **Figure 2B**) appeared in the contralateral visual hemifield relative to the recording hemisphere (**Figure 2A**, ‘peri. rings’). Fixation on FR was required throughout trial. Subsequently, an attention cue (red filled circle) appeared for 0.4 s on one of the three rings, indicating the target ring for that trial (the (attend) Up, Down or FR conditions). After a random waiting period (wait1, 1.7-4.8 s), in 50% of trials (short trials), the color of the target ring briefly changed to red (target change), requiring the monkeys to release the lever within 0.8 s from the change (lever release). In the remaining 50% (long trials), the wait1 period ended with a color change in one of the two non-target rings (catch change), to which the monkey was prohibited from responding. The monkeys then waited an additional 1.0-3.2 s (wait2 period) before the target change. In the single-task attention trials (i.e., trials without MGS-task insertion), a correct lever release was rewarded with a drop of juice, either immediately (50% of trials) or after an additional 0.4–1.0 s of central fixation (50%; randomly selected per trial). The two reward timing conditions were introduced to examine the reproducibility of a prior study^11^ concerning the role of the monkey FPC in decision monitoring (see Experiment 5 below).

In two-thirds (66.7%) of randomly selected attention task trials, the MGS task was inserted partway through the trial by a memory cue presentation (0.25 s, **Figure 2D**) during either the wait1 or wait2 period at one of six far-peripheral locations (**Figure 2C**), creating dual-task trials (dual MGS task, ‘DMT’). MGS insertion was equally balanced between short and long attention task trials, with each type comprising 33.3% of all attention task trials. The memory cue appeared at random intervals after offset of the attention cue, between 1.0 to 4.1 s in short trials and 1.0 to 5.0 s in long trials of the attention task. The monkeys were required to memorize this location while continuing with the attention task. After the attention task was complete (lever release) and a subsequent follow-up fixation (0.4-1.0 s), all rings disappeared, and six small placeholders appeared (saccade go-signal). The monkeys then made a saccade (<0.5 s) to the placeholder where the memory cue had been presented and maintained gaze for 0.25 or 0.6 s (blocked) before receiving a reward. Thus, in DMT, the identical MGS task was performed concurrently with the three attention task conditions (Up, Down, and FR); hereafter referred to as DMT-up, DMT-down, and DMT-FR (**Figure 2E**). As a control, in separate blocks, the monkeys performed the MGS task alone (single MGS task, SMT), which had the same time course as in DMT, with all attention task events scheduled but executed without any physical changes. By comparing neural activity between SMT and DMT throughout the LPFC, we aimed to reveal previously unknown subregional differences in neuronal contributions to dual-task cognitive orchestration. **Methods** describes the MGS-insertion rule; **Figure S1** shows example DMT and SMT trial sequences.

During daily recording sessions (38 for monkey T and 22 for monkey K), a DMT block was performed between two SMT blocks. Monkey K also completed 15 ‘frequent task-switching sessions,’ where SMT and DMT blocks alternated frequently (8 ± 1.2 times per session; every 40–50 correct trials). Unless otherwise noted, all analyses were performed using data from all sessions. An analysis of attention task performance (**Figures 2F** and **2G**, paired-permutation test; all *p*-values corrected) showed that in both monkeys, response time (RT) primarily reflected task difficulty, and the Up and Down conditions (detection of target changes in the periphery) were significantly more difficult than the FR condition (foveal detection). In MGS (**Figure 2H**), monkey T (left panel) showed a significant performance decrement in DMT-down (*p* = 6.0 × 10^−4^) and a marginally significant decrement in DMT-up (*p* = 0.06) compared to SMT. In monkey K (right), although no significant performance decrement was observed, when trials with a longer memory delay period (> 3.0 s, dashed red line graph) were analyzed separately, a marginally significant decrement in DMT-down (*p* = 0.08) was observed. In the combined data from the two monkeys (*n* = 75), dual-task performance decrement was highly significant in DMT-down (*p* = 4.0 × 10^−4^) and marginally significant in DMT-up (*p* = 0.07) (all *p*-values corrected). These results indicate that adding the attention task increased task complexity and coordination demands compared to SMT, consistent with our previous reports^16,24^.

To examine how dual-task performance affected MGS activity, we conducted a two-way ANOVA with the following factors: (cue) location (six levels) and task (SMT or DMT) for each task period in each neuron. **Figure 3A** shows a representative dorsal pos-PFC neuron that exhibited a significant reduction in location selectivity during DMT compared to SMT in the delay period (grey shaded area, 0.4 – 1.0 s from memory cue onset; interaction, *F*_5,314_ = 15.0, *p* = 3.9 × 10^−13^). This neuronal dual-task interference effect^16^ likely reflects an allocation of processing resources away from the MGS and toward the attention task. However, following completion of the attention task (lever release, ‘lever rel.’ in DMT), this neuron regained its suppressed location selectivity toward the saccadic response, even surpassing that in SMT (memory reactivation^16,25^), suggesting a reallocation of resources back to the MGS. This activity likely reflected an adaptive distribution of cognitive resources between the two tasks.

**Figure 3.**
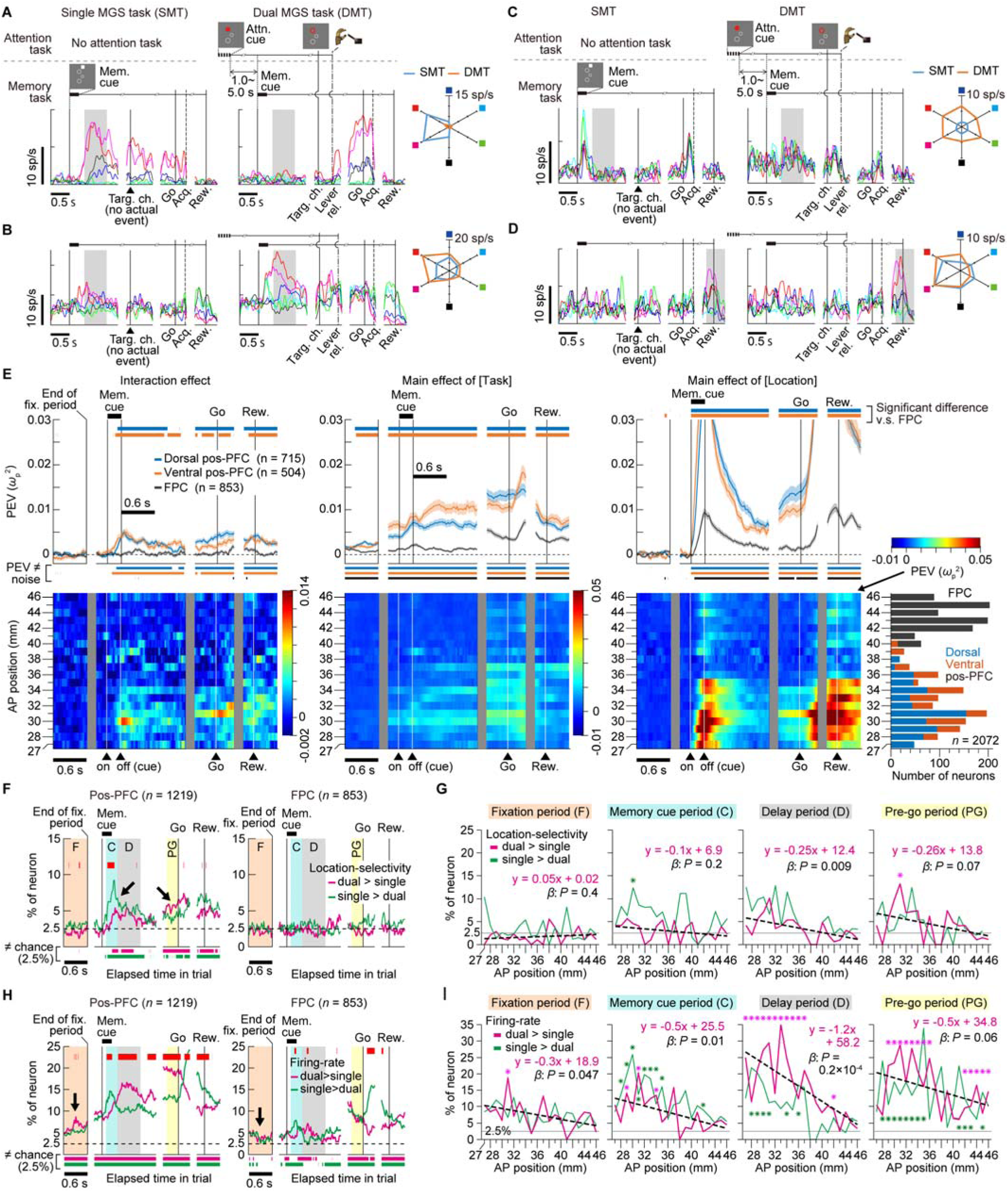
Neural activity related to cognitive orchestration across the anteroposterior LPFC. (**A**) Activity of a representative dorsal pos-PFC neuron during the single MGS task (SMT, left) and dual MGS task (DMT, middle). Activity is aligned to the onset of memory cue (mem. cue), target change (targ. ch.), go-signal (go), and reward (rew.). Each spike-density function color corresponds to the memory cue location indicated in the polar plot (right panel) which shows the mean firing rate during the delay period (grey-shaded area) for each cue location. Vertical dashed lines indicate the mean lever-release time (lever rel., DMT only) and mean saccadic target acquisition time (acq.). (**B** to **D**) Activity of a representative dorsal pos-PFC neuron (**B**); ventral pos-PFC neuron (**C**); and FPC neuron (**D**). The polar plot in **D** shows the peri-reward period activity (gray-shaded area). (**E**) Upper row: Time course of population PEV for interaction term (left), and main effects of task (middle) and location (right) during MGS. Activity is aligned to the end of the fixation (fix.) period, the onset of memory cue, go-signal, and reward. The fixation period is the initial fixation interval at the beginning of the attention task component (0.6–1 s; Figure 2D). Shaded areas indicate S.E.M. Lower horizontal bars mark time periods of significant PEV (i.e., ≠ noise) in each recording area (one-sample permutation test, *p* < 0.05; FDR-corrected). Upper horizontal bars mark time periods when PEV in dorsal (cyan) or ventral (orange) pos-PFC was significantly different from that in the FPC (two-sample permutation test, FDR-corrected). Bottom row: Time course of population PEV at 1-mm increments along the AP axis. Rightmost panel shows the number of recorded neurons in each 1-mm segment. Bin width = 200 ms (slid in 20 ms). All analyses excluded neural responses evoked by attention-task events (**Methods**). (**F**) Time course of the percentages of location-selectivity_dual>single_ neurons (magenta) and location-selectivity_single>dual_ neurons (green). Upper red horizontal bar indicates time periods when the percentage differed significantly between the two neuron types. Lower horizontal bars indicate time periods when the percentage of each neuron type significantly differed from chance (2.5%) (Fisher’s exact test, *p* < 0.05; FDR-corrected). (**G**) Changes in the percentages of the location-selectivity_dual>single_ (magenta) and location-selectivity_single>dual_ (green) neurons along the AP locations during the fixation (orange), memory cue (cyan), delay (gray) and pre-go (yellow) periods. The durations of these periods are shown in **F**. Regression lines and equations indicate the rate of change in the percentage of location-selectivity_dual>single_ neurons as a function of AP position. Asterisks indicate percentages significantly different from chance (2.5%) (FDR-corrected). (**H** and **I**) Same as in **F** and **G**, but for the firing-rate_dual>single_ (magenta) and firing-rate_singlel>dual_ neurons (green).

The dorsal pos-PFC neuron in **Figure 3B** showed the opposite interaction effect during the delay period (location selectivity, DMT > SMT; *F*_5,344_ = 11.9, *p* = 1.3 × 10^−10^), suggesting selective participation in mnemonic processing under DMT. The ventral pos-PFC neuron in **Figure 3C** exhibited a significant main effect of task during the delay period (*F*_1,241_ = 166.21, *P* = 2.8 × 10^−29^) with a higher overall firing rate across cue locations in DMT than in SMT. These two DMT-enhanced activities likely contributed to dual-task-specific processes. Additionally, many FPC neurons (**Figure 3D**) exhibited a significant effect of location only during the peri-reward period (grey shaded area, −0.2 to 0.3 s from reward, simple-effect *p* < 0.0002 for both SMT and DMT), consistent with prior findings in a spatial cued-strategy task^11^.

We calculated the time course of population-averaged partial omega-squared proportion of explained variance (PEV) for each factor (**Figure 3E**). A brain area that is critical for dual-task processing should show significant PEV in the interaction term and/or the factor task. However, for both, the FPC’s PEV was significantly lower than that of the dorsal and ventral pos-PFC throughout the trial (top row, two leftmost panels), indicating that FPC activity made a significantly weaker distinction between single- and dual-task conditions than the pos-PFC. A segment-by-segment analysis along the AP axis (bottom row) confirmed that the distinction was strongest at AP levels corresponding to the posterior third of the principal sulcus (AP 27 – 33 mm, spanning both intra- and extra-sulcal regions), moderate in the middle third (AP 33 – 39 mm), and minimal in the anterior third and beyond (AP > 39 mm).

### Dual-task-specific information processing emerges primarily in the posterior-to-mid LPFC

Dual-task-specific information processing likely involves two neural populations. The first is comprised of neurons exhibiting a significant interaction effect, with stronger location selectivity in DMT compared to SMT (**Figure 3B**, hereafter ‘location-selectivity_dual>single_ neurons’). This activity can be considered ‘content-specific recruitment,’ covering increased demand on task-content processing under DMT. The second population is comprised of neurons exhibiting a significant main effect of task, with higher overall firing rates in DMT than in SMT (**Figure 3C**, ‘firing-rate_dual>single_ neurons’). This activity reflects ‘load-indexing recruitment’, scaling with overall task difficulty or complexity, and likely triggering dual-task-specific processing in other neural populations. To determine when and where these recruitment patterns occurred, we calculated the time course of the percentages of these neurons among all recorded neurons in each region. For comparison, we also calculated the percentages of significant single-task-preferring neurons (location-selectivity_single>dual_ neurons and firing-rate_single>dual_ neurons; **Figures 3F** and **3H**, green curves). Hereafter, we collapsed data from dorsal and ventral pos-PFC.

The results showed that in the pos-PFC (**Figure 3F**, left panel), but not in the FPC (right), the percentage of location-selectivity_dual>single_ neurons (magenta) significantly exceeded chance (2.5%) during the memory cue and delay periods (cyan and gray shaded areas), as well as throughout the trial, indicating that an add-on information-processing mechanism that operates exclusively during multitasking exists only in the pos-PFC. For load-indexing recruitment (**Figure 3H**), only in the pos-PFC (left panel) did the dual-task-preferring firing-rate_dual>single_ neurons significantly outnumbered the firing-rate_single>dual_ neurons throughout the delay and pre-go periods, indicating that only the pos-PFC exhibited higher global firing rates in response to an increasing number of competing goals. To our knowledge, these two types of dual-task-specific activity constitute the first evidence of prefrontal neuronal recruitment during cognitive orchestration. They may correspond to the selective BOLD-signal increase in human anterior LPFC during multitasking^26,27^.

Next, across each 1-mm segment of recording location, we examined how the percentages of location-selectivity_dual>single_ neuron and firing-rate_dual>single_ neurons changed along the AP axis during the fixation, memory cue, delay and the pre-go periods (magenta plots in **Figures 3G** and **3I**; see color-shaded areas, ‘F’, ‘C’, ‘D’ and ‘PG’ in **Figure 3F** for the analysis time window). We found that, except for the fixation and memory cue periods in location-selectivity_dual>single_ neurons (**Figure 3G**, two leftmost panels, magenta), both the percentages of location-selectivity_dual>single_ and firing-rate_dual>single_ neurons showed a significant or marginally significant anteriorly decreasing trend (for all β, *p* < 0.07), with the most anterior locations predominantly exhibiting non-significant percentages. This again suggests that the FPC plays minimal roles in cognitive orchestration. Notably, only in the pos-PFC (**Figure 3H**, left), starting from the fixation period, the percentage of firing-rate_dual>single_ neuron rose significantly above both chance and the percentage of firing-rate_single>dual_ neurons (arrow), suggesting that, even from the preparatory period, only the pos-PFC maintained heightened readiness for the anticipated higher cognitive demand in DMT.

Interestingly, the results in **Figure 3F** (left panel) reveal another dual-task specific phenomenon exclusively in the pos-PFC: an adaptive resource allocation between the two tasks. Consider again the representative neuron in **Figure 3A**. During the memory cue and delay periods, this neuron exhibited single-task-dominant selectivity (location-selectivity_single>dual_) due to dual-task interference. At the population level (**Figure 3F**, left panel), we observed this pattern as a higher proportion of location-selectivity_single>dual_ than location-selectivity_dual>single_ neurons during the same periods (green curve exceeding magenta, left diagonal arrow), suggesting a net diversion of resources from the MGS to the concurrent attention task. However, approaching the go time (after attention-task completion), the same representative neuron (**Figure 3A**) shifted to dual-task-dominant selectivity (location-selectivity_dual>single_), displaying the memory reactivation phenomenon. This shift was again evident in the pos-PFC population, reflected by an increasing proportion of location-selectivity_dual>single_ neurons near go time (**Figure 3F**, left panel; magenta exceeding green, right diagonal arrow), suggesting that resources were reallocated back to MGS. This crossover, from initial emphasis on the attention task (green exceeding magenta) to subsequent emphasis on MGS (magenta exceeding green), illustrates dynamic resource allocation between the two competing tasks in the pos-PFC, a phenomenon absent in the FPC.

Together, the results of Experiment 1 indicate that, contrary to the dominant view in human studies^3^ and a previous monkey lesion study^8^, the simultaneous performance of multiple cognitive tasks did not elicit selective activations in the monkey FPC. Instead, the orchestration process appears to be predominantly supported by the posterior-to-mid LPFC (up to approximately AP 34 mm), where significant percentages of dual-task specific activity was observed. Results were essentially unchanged when we performed the same set of analyses on neurons recorded during the aforementioned frequent task-switching sessions which controlled for potential firing-rate drift due to the block-design session structure with long blocks (**Figure S2** and **Supplementary Text**)

### Experiment 2: Testing the cognitive orchestration hypothesis in a serial multimodal dual task

To rule out task-specific effects, we used another dual-task paradigm in which secondary tasks were inserted during the ITI of an object few-shot learning (FsL) task. This structure, involving the serial presentation of tasks from different modalities, parallels that of a prior FPC lesion study^8^. In the FsL task, monkeys worked through a series of *problems* by searching for and exploiting a rewarded object stimulus (S^+^) among three objects (**Figure 4A**; S^+^, S_1_^−^ and S_2_^−^; size, 3.8° on a side). The identities of S^+^, S_1_^−^ and S_2_^−^ remained the same within each problem which lasted until the completion of 8 – 9 correct trials. Each problem ended with a 10–15 s blank interval before a new problem began, introducing three novel objects. Two different spatial configurations were used for object presentation in alternate trials, with the three objects placed randomly within each configuration (**Figure 4B**).

**Figure 4.**
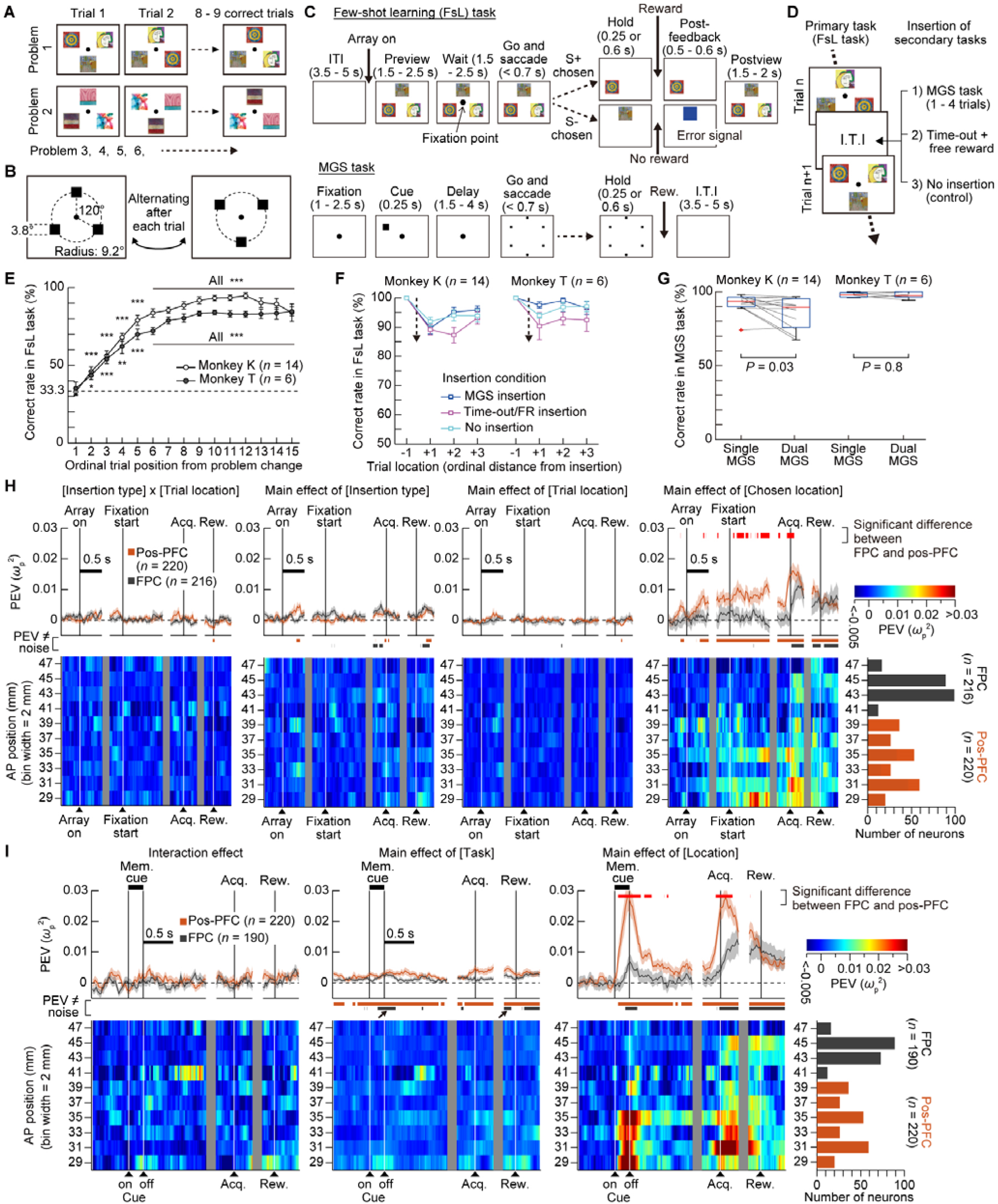
Neural activity during the serial multimodal dual task across the anteroposterior LPFC. (**A**) Progression of the few-shot learning (FsL) task. Monkeys K and T performed an average of 36.4 and 37.7 problems per session. (**B**) Two spatial configurations for object presentation in the FsL task, alternating each trial. (**C**) Event sequences of the FsL (upper row) and MGS (bottom row) tasks. (**D**) Insertion of a secondary task during the ITI of the FsL task. (**E**) Behavioral performance in the FsL task (one-sample *t*-test, FDR-corrected). (**F**) Differential effects of three secondary-task insertions on FsL performance. Dotted arrows indicate the timing of insertion. (**G**) Comparison of correct response rates between single-task and dual-task MGS. (**H**) Time course of population PEV (mean ± s.e.m.) for the four key terms in the FsL, aligned at the start of the preview period (‘array on’) and wait period (fixation start), and the timing of target acquisition (acq.) and reward (rew.). Upper row: Red horizontal bars mark periods of a significant difference in PEV between the FPC (gray curve) and pos-PFC (orange curve) (FDR-corrected). Lower colored horizontal bars mark periods when PEV was significantly different from noise in each recording area (FDR-corrected). Bottom row: Time course of PEV in each 2-mm segment along the AP axis, with the rightmost panel showing the number of recorded neurons. (**I**) Same as in **H**, but for the comparison between the single-task and dual-task MGS. Note that due to a program error in the first session for monkey T, the cue location of the dual-task MGS was not recorded, and therefore 26 FPC neurons were removed from this analysis.

On each trial (**Figure 4C**, top), after an intertrial interval (ITI, 3.5 – 5.0 s), the monkey was presented with an object array (‘array on’) and freely viewed it (‘preview’, 1.5 − 2.5 s). Then, a central fixation point (FP) was presented, and the monkeys were required to fixate it. After a variable wait period (‘wait’, 1.5–2.5 s), the disappearance of the FP prompted the monkeys to choose one of the three objects by saccade (‘go and saccade’), with unchosen objects disappeared upon choice. The monkeys then fixated on the chosen object for 0.25 or 0.6 s (‘hold’). Selection of S^+^ led to a reward, while selection of S_1_^−^ or S_2_^−^ resulted in an error feedback (a blue patch replacing the chosen object for 0.5-0.6 s; ‘post-feedback’). Then the monkeys were again presented with and freely viewed the original display (‘post-view’). During ITI, the monkeys were required to maintain the memory of S^+^.

On some trials, one of three secondary-task conditions was randomly inserted during the ITI: MGS insertion, time-out/free-reward insertion (’time-out/FR’) or no insertion (**Figure 4D**). In the MGS insertion condition, 1-4 successive MGS trials (**Figure 4C**, bottom) were inserted. In the time-out/FR condition, a blank screen was shown for 40-50 s, during which 1-4 free rewards were delivered at random intervals, triggering attention-drawing events (as in the insertion of “reward consumption” task in the previous FPC lesion study^8^). In the no-insertion condition, nothing was inserted (i.e., single-task control condition). In addition to this comparison between single- and dual-task FsL, to complement Experiment 1, we introduced a block of single MGS task before and after the FsL block (70 – 100 trials each) and compared neural activity between the single-task MGS and the MGS inserted during FsL performance (dual-task MGS).

In the FsL task, Monkeys K and T showed significantly above chance (33%) correct rate from the second trial after problem change, indicating few-shot learning of novel stimulus-reward (S-R) associations in each problem (**Figure 4E**). To examine the effect of secondary-task insertion on FsL performance, we analyzed correct rate changes before and after insertion. Since our focus was how the memory of S^+^ was affected by secondary tasks, we only analyzed cases where the trial immediately before insertion was correct. Different insertion conditions led to varying performance decrements in the post-insertion FsL trials (**Figure 4F**). In both monkeys, the time-out/FR insertion caused the largest performance decrement, while the MGS insertion had a similar effect to the no-insertion. This differential effect of insertion condition was marginally significant in monkey K (two-way ANOVA: interaction, *F*_6,78_ = 2.0, *p* = 0.08; main effect of insertion type, *F*_2,26_ = 2.3, *P* = 0.12; trial location*, F*_3,39_ = 19.3, *p* = 8.0 × 10^−8^). In monkey T, the interaction and the main effect of insertion type were not significant (*F*_6,30_ = 1.5, *p* = 0.22, and *F*_2,10_ = 2.8, *p* = 0.11, respectively), likely due to the small number of sessions (*n* = 6). However, in combined data for both monkeys (*n* = 20), both effects reached significance (interaction, *F*_6,114_ = 2.2, *p* = 0.049; insertion type, *F*_2,38_ = 4.7, *p* = 0.01). Between the single- and dual-task MGS (**Figure 4G**), only monkey K showed a significant dual-task performance decrement (*p* = 0.03; monkey T, *p* = 0.8). However, in combined data (*n* = 20), significant dual-task performance decrement was observed (*p* = 0.04).

We recorded activity from 216 FPC and 220 pos-PFC neurons (**Figure 1I**). To assess dual-task-specific representations, we computed the time course of population-averaged PEV in the FsL task, using a three-way ANOVA with factors: insertion type (3 levels), trial location (pre- or post-insertion), and chosen location (6 levels) (**Figure 4H**). If the monkey FPC is involved in dual-task-specific processing, we would expect a significant interaction (insertion type × trial location) and/or a simple main effect of insertion type in the post-insertion trials where the task diverged into a single or serial dual-task format. However, in both the FPC and pos-PFC, significant PEV was absent in both the above interaction term (**Figure 4H**, leftmost panel) and the simple main effect (**Figure S3A**). The only consistently significant PEV was for chosen location (**Figure 4H**, rightmost panel), a task-irrelevant spatial factor, with significant PEV in the FPC emerging only after target acquisition (acq.). These findings indicate that in this particular dual-task, both FPC and pos-PFC activity play a minimal role. These observations were further validated by the PEV across 2-mm recording segments (bottom row). Calculating the percentage of significant neurons yielded nearly identical results (**Figure S3B**).

Next, to compare neural activity between the single- and dual-task MGS, we calculated the population PEV using a two-way ANOVA with factors task (single vs. dual) and location (6 levels) (**Figure 4I**). The absence of significant interaction (leftmost panel) indicates that location selectivity did not differ between the single and dual MGS tasks in both FPC and pos-PFC. For the factor task, the FPC showed significant PEV only during two brief epochs (140-440 ms from cue onset and −200 to −80 ms from reward; arrows in middle panel), whereas the pos-PFC exhibited significant PEV throughout the trial, as in Experiment 1. These results indicate that only the pos-PFC discriminated between the single- and dual-task MGS.

We applied the same analysis used in Experiment 1 (**Figure 3H**) to identify the source of this effect (**Figure S3D**). The results showed that, in the pos-PFC, the dual-task-preferring firing-rate_dual>single_ neurons again outnumbered firing-rate_single>dual_ neurons throughout the task. This indicates that, as in Experiment 1, the pos-PFC exhibited higher overall firing rates with an increasing number of competing goals (load-indexing recruitment), despite different combinations of task modality (space only in Experiment 1 vs. space and object in Experiment 2) and task structure (parallel vs. serial dual-task). Finally, the PEV for location (**Figure 4I**, rightmost panel) resembled that in Experiment 1. Calculating the percentage of significant neurons yielded comparable results (**Figure S3C**).

Together, the results from Experiments 1 and 2 indicate that, despite the vastly different task settings, only the pos-PFC adaptively modulates activity from single- to dual-task conditions. This suggests that monkey FPC is unlikely to play essential roles in dual-task-specific processes such as task coordination^26,27^ and resource allocation^8,15^. In the pos-PFC, the observed neural dual-task effect was smaller in Experiment 2 than in Experiment 1. This is likely because, in Experiment 2, the serial execution of tasks with different modalities minimized neuronal competition between the tasks and reduced the need for dual-task processing^28^. In contrast, Experiment 1 required the simultaneous execution of two spatial tasks, which likely resulted in more severe competition for processing resources^16^ and necessitated mediation by orchestration processes.

### Experiment 3: Testing the rapid novel learning hypothesis of FPC

In a previous study, an FPC lesion impaired rapid learning of novel object values in a task similar to the present FsL task (Figure 2D in Ref. 7, *p* = 0.048, one-tailed *t*-test). Nougaret et al. (2024)^13^ reported selective modulation of FPC activity during initial novel learning. In their Object-in-Place (OiP) task, where monkeys searched for and exploited a small rewarded target embedded in novel full-screen scenes, a full-sample analysis of all 399 recorded FPC neurons (i.e., no prescreening; Supplementary Figure 2 in Ref. *13*) found that 6.0% (24/399) responded more to novel than familiar scenes (novelty-preferring cells), while 1.8% (7/399) exhibited the opposite pattern (familiarity-preferring cells). Both studies suggested that the FPC plays a leading role in the initial stage of novel learning.

However, to comprehensively assess FPC’s contribution, it is essential to compare it with other regions involved in novel learning, including the posterior LPFC^17,20,29^. Thus, we continued recording in the FsL task (**Figure 1J**) with slight modifications (**Figure 5A**, modified FsL task): (i) omitting the preview and postview periods, and secondary-task insertions, and (ii) requiring monkeys to maintain central fixation from 1.0–1.5 s before array onset until the go-signal. In separate blocks, we recorded neural activity during the MGS task using the same parameters as in Experiment 2, occasionally delivering free rewards during the ITI

**Figure 5.**
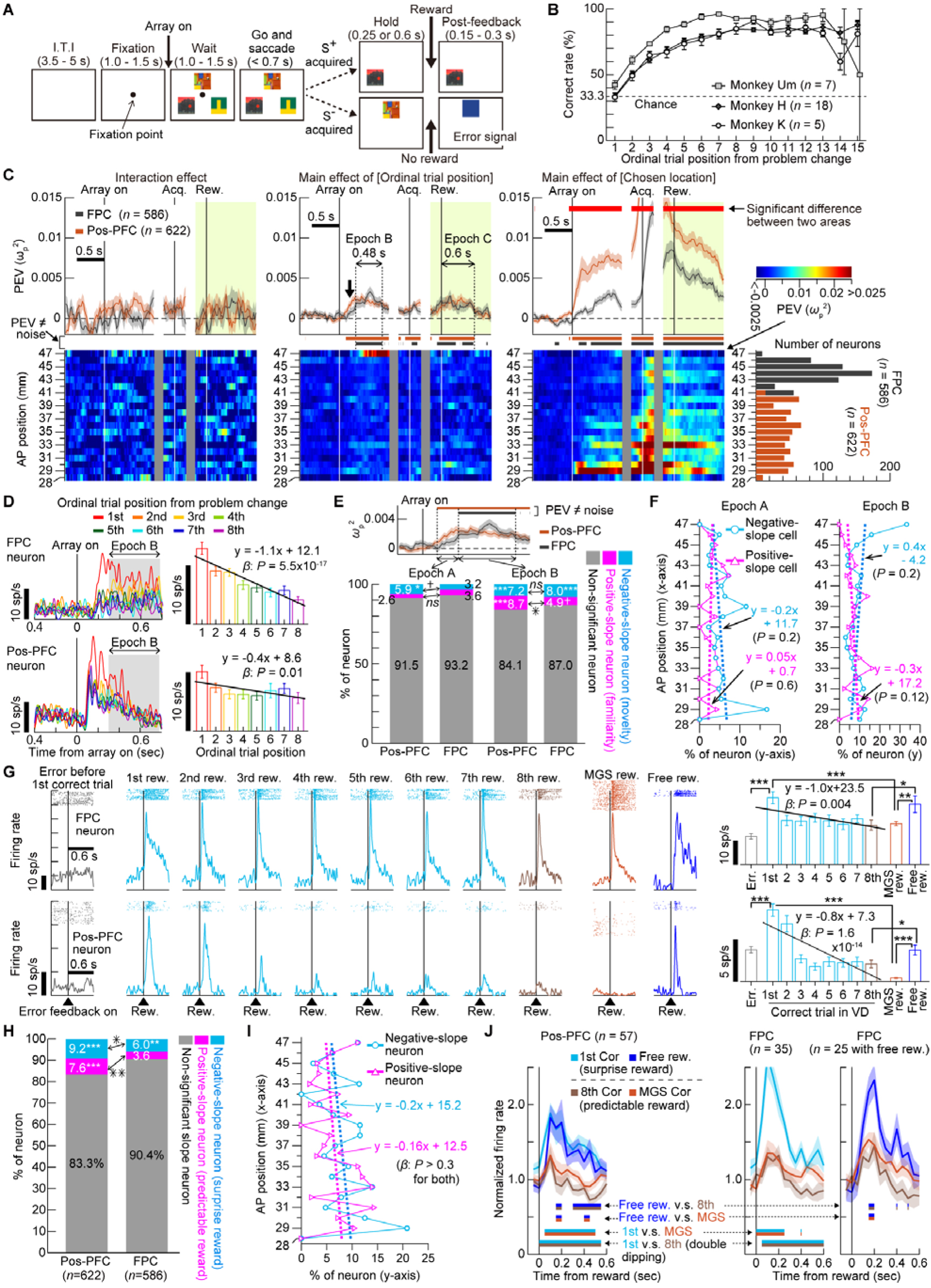
Neural activity related to novel learning across the anteroposterior LPFC. (**A**) Trial event sequence of the modified FsL task. Each problem lasted until the completion of 8 correct trials. (**B**) Behavioral performance. Monkeys K, H, and Um completed an average of 47.0, 42.1 and 32.7 problems per session. (**C**) Time course of population PEV (mean ± s.e.m.) in the FPC (gray) and pos-PFC (orange), aligned to the start of the wait period (array on), saccadic target acquisition (acq.) and reward (rew.). Other conventions as in Figure 4H. (**D**) Activity of a representative negative-slope neuron in the FPC (top) and pos-PFC (bottom) across trials 1 to 8 after problem change (regardless of correct/error outcome; fixation-break error trials excluded). Right bar graphs illustrate the mean firing rate in Epoch B (gray shaded area) and an OLS regression result. (**E**) Percentage of negative- (cyan) and positive-slope neurons (magenta) in the pos-PFC and FPC for Epochs A (left) and B (right). The inset PEV plot enlarges the region near ‘array on’ in **C** (second panel from left). *P*-values are FDR-corrected for six comparisons per epoch. (**F**) Percentage of negative- (cyan) and positive-slope neurons (magenta) at each 1-mm AP location. For clarity, the x-value is plotted on the vertical axis and the y-value on the horizontal axis. (**G**) Epoch C (post-reward period): Activity of a representative negative-slope neuron in FPC (top) and pos-PFC (bottom) during the 1st to 8th correct trial after problem change. Also shown are responses during error trials, MGS reward, and free reward. The rightmost panel summarizes the mean firing rate for each trial type in Epoch C with OLS regression results (err. = error). (**H**) Same as in **E**, but for Epoch C. (**I**) Same as in **F**, but for Epoch C. (**J**) Comparison of normalized population activity of negative-slope neurons between surprise rewards (1st correct trial and free reward, cold colors) and predictable rewards (8th and MGS correct trials, warm colors) in the pos-PFC (left) and the FPC (right). Bin width 50 ms. Shaded areas indicate within-subject SEM. Lower horizontal bars mark periods of a significant difference between the two conditions as specified. For FPC data, due to a timestamp error in the first two sessions for monkey T, free reward responses were not recorded for 10 negative-slope neurons. Thus, comparisons of free vs. 8th rewards and free vs. MGS rewards (rightmost panel) were performed separately for the remaining 25 neurons.

In the modified FsL task, all monkeys (K, H and Um) exhibited significantly above-chance (33%) performance from the second trial after problem change, indicating rapid novel learning (**Figure 5B**). Combining data from Experiments 2 and 3, we analyzed 586 FPC and 622 pos-PFC neurons. For each region, we calculated the time course of population PEV using a two-way ANOVA with factors: ordinal trial position from problem change (8 levels: 1st to 7th, and ≥8th trials combined) and chosen location (6 levels) (**Figure 5C**). Except for the peri-reward period (green shaded areas, top panels), both correct and error trials were analyzed (fixation-break error trials excluded). For the peri-reward period, only correct trials were analyzed, with the factor ordinal trial position representing the number of *correct* trials from problem change (1st through 8th). Brain regions crucial for rapid novel learning should exhibit differential activations as a function of the ordinal trial position from problem change (i.e., significant PEV for the interaction and/or the factor ordinal trial position).

In both FPC and pos-PFC, the interaction remained nonsignificant throughout the trial (**Figure 5C**, left panel, permutation test, FDR-corrected *p* > 0.05), indicating that chosen location selectivity did not change with novel S-R learning progression. However, for the factor ordinal trial position, two relatively long intervals showed significant PEV in both regions (middle panel: ‘Epoch B,’ 300 – 780 ms from array onset; ‘Epoch C,’ 0 – 600 ms from reward). This indicates that overall firing rates (averaged across all chosen locations and objects) were significantly modulated as learning progressed, suggesting that both regions distinguished between exploration and exploitation phases.

In Epoch B, to illustrate these population-level effects at the single-neuron level, **Figure 5D** shows the activity of representative FPC (top panels) and pos-PFC neurons (bottom) in the modified FsL task (**Figure 5A**). On the first trial of each problem, both neurons exhibited the strongest response at array onset, followed by a significant reduction in subsequent trials. Both neurons showed significantly negative slope in an ordinary least squares regression (OLSR) (right panels) with ordinal trial position as the explanatory variable and trial-by-trial firing rates in Epoch B as the response variable, which was the sole criterion for classifying a neuron as a ‘negative-slope neuron’. Because novel objects were introduced in each problem, these activities represent object-invariant signals of novel behavioral contexts. We also identified neurons with opposite preference, those with significantly positive slopes in the OLSR (positive-slope neuron, **Figure S4A** and **S4B**), which signaled object-invariant occurrences of familiar contexts.

### Posterior-to-mid LPFC detects novel contexts earlier than FPC

One notable trend before Epoch B (**Figure 5C**, middle panel) is that, following array onset, a significant PEV increase in the pos-PFC preceded that in the FPC by 180 ms (arrow, ‘Epoch A’; see **Figure 5E** for an enlarged view). This delayed elevation suggests that the FPC may play only a subordinate role in detecting novel contexts compared to the pos-PFC, a view not tested in previous studies^7,13^. To address this, we examined whether, for each epoch, the percentage of negative- and positive-slope neurons in each area exceeded chance (2.5%) and differed between areas (**Figure 5E**; Fisher’s exact test, *p*-values FDR-corrected for six comparisons per epoch). In Epoch A (two left bars), only the percentage of negative-slope neurons in the pos-PFC (cyan) was significantly above chance (5.9%, *p* = 0.03). The other three values were non-significant (positive-slope neuron in pos-PFC, 2.6%; negative- and positive-slope neuron in FPC, 3.2% and 3.6% respectively; *p* > 0.6 for all). Between areas, the percentages of negative-slope neurons in the pos-PFC was marginally significantly higher than that in the FPC (*p* = 0.08), whereas the percentages of positive-slope neurons did not differ (*p* = 0.6). This indicates that the pos-PFC exhibited a greater proportion of negative-slope neurons at an earlier time point than the FPC.

In Epoch B (**Figure 5E**, two rightmost bars), all percentages were significantly or marginally significantly above chance (pos-PFC: 7.2% and 8.7%; FPC: 8.0% and 4.9% for negative- and positive-slope neurons, respectively; *p* < 0.054 for all four comparisons). Between the areas, pos-PFC had a significantly higher percentage of positive-slope neurons than FPC (*p* = 0.02), but no such difference was observed for negative-slope neurons (*p* = 0.67). This indicates that the FPC’s contribution to novel learning catches up to the pos-PFC only at Epoch B. Notably, the observed percentage of negative-slope neurons in Epoch B in the FPC (8.0%, 47/586 neurons; rightmost bar) closely matched (*p* = 0.26) that of the novelty-preferring FPC cells in Nougaret et al.^13^ (6.0%; 24/399) during a comparable task period in their OiP task (200– 600 ms from scene onset). In the present study, by directly comparing pos-PFC and FPC activity in the same animals and task, we showed that in novel few-shot learning, the FPC follows, rather than precedes, pos-PFC activation. Repeating the analysis for each 1-mm AP location (**Figure 5F**) revealed no evidence that the percentages of negative- or positive-slope neurons in any FPC segments exceeded chance (2.5%) during either Epoch A or B (*p* > 0.1 in all 20 comparisons at each AP location, Fisher’s exact test, FDR-corrected). The seemingly high value (33%) of negative-slope neurons at AP 47-mm bin in Epoch B is likely an outlier due to the small sample size (*n* = 9) (vs. chance, *p* = 0.59).

One concern is that the present analysis included the data from both fixation (Experiment 3) and non-fixation (Experiment 2) conditions before and after array onset potentially confounding neural activity with eye movements near array onset. This issue also applies to the prior FPC study^13^, where fixation was not required at all during the OiP task. To address this, we repeated the analyses using Experiment 3 data. The results (**Figure S5** and **Supplementary Text**) were essentially unchanged.

### Posterior-to-mid LPFC more strongly encodes reward prediction errors than FPC

In Epoch C (0–600 ms after reward; **Figure 5C**, middle panel), we newly identified a reward-related type of negative-slope neuron in both FPC and pos-PFC (representative neurons, **Figure 5G**). Their activity decreased significantly from the 1st to 8th reward after problem change, with a significantly negative slope in OLSR (rightmost panels). Additionally, in both neurons, (i) delivery of free (surprise) reward elicited significantly greater activation than the 8th (predictable) reward (rightmost panels; paired permutation test, *p* < 0.03 for both neurons, FDR-corrected for four comparisons per neuron); and (ii) responses to both the 1st and free rewards (both surprise) were significantly greater than those to the MGS (predictable) reward (*p* < 0.004). These negative slope (reward) neurons signal the occurrence of unexpected rewards and exhibit characteristics of a reward prediction error (RPE) signal which promotes learning by triggering updating of prior S-R relations^30^. We also observed positive-slope neurons (**Figure S4C**), which likely signal the delivery of predictable rewards. A brain area more involved in novel learning would show a higher proportion of negative-slope neurons. Indeed, the pos-PFC contained a significantly higher percentage of both negative-slope and positive-slope neurons (9.2% and 7.6%, respectively, **Figure 5H**) than the FPC (6.0% and 3.6%) (Fisher’s exact test, *p* = 0.047 and *p* = 0.005, FDR-corrected as in **Figure 5E**). Across each 1 mm segment of the recording area, both neuron types showed an anteriorly decreasing trend, though not significant (**Figure 5I**, *p* > 0.3 for both). Finally, we compared normalized population-averaged firing rate of all negative-slope neurons in response to surprise rewards (1st and free, cold colors) versus predictable rewards (8th and MGS, warm colors; **Figure 5J**). The neural responses to both types of surprise rewards were highly similar to each other and significantly exceeded those to predictable rewards in both the pos-PFC and FPC (paired-permutation test, FDR-corrected), suggesting that population-activity of negative slope (reward) neurons in both areas represents RPE signal.

Taken together, these findings indicate that, while FPC activities at array onset and after reward distinguish exploration from exploitation during novel learning, this distinction is significantly weaker than in the pos-PFC: (i) it emerges later (array-onset response in Epoch B) and (ii) involves fewer neurons (reward response in Epoch C), suggesting a subordinate role for the FPC in novel learning. Notably, even in pos-PFC, negative-slope neurons comprised fewer than 10% of recorded neurons at both array onset and post-reward. Given that the ventral corticostriatal circuit, including OFC, primarily drives value-based object learning^23,31^, the pos-PFC itself may have only a modest role.

### Experiment 4: Testing the rapid learning hypothesis in a small-set FsL task

In Experiment 3, the introduction of novel objects in each problem prevented the use of chosen-object identity as an experimental factor, so we could not directly test whether FPC neurons encode valuable objects from the earliest timing in novel learning. To address this, we further modified the FsL task to present only five objects per session (the ‘small-set FsL task’; **Figure 6A**). The task (**Figure 6B**) proceeded as in Experiment 3, except that: (i) each problem continued until the completion of 5 or 6 correct trials; (ii) in each problem, two objects (S^+^ and S^−^) were randomly selected from the five session-specific objects (**Figure 6A**, bottom right); and (iii) on each trial, S^+^ and S^−^ were placed at two of three possible locations (90°, 210°, and 330°; **Figure 6A**, top right). In separate blocks, we recorded neural activity during the MGS task (**Methods**). We recorded activity from 192 FPC and 213 pos-PFC neurons in monkeys K and H (**Figure 1K** and **Table S1**)

**Figure 6.**
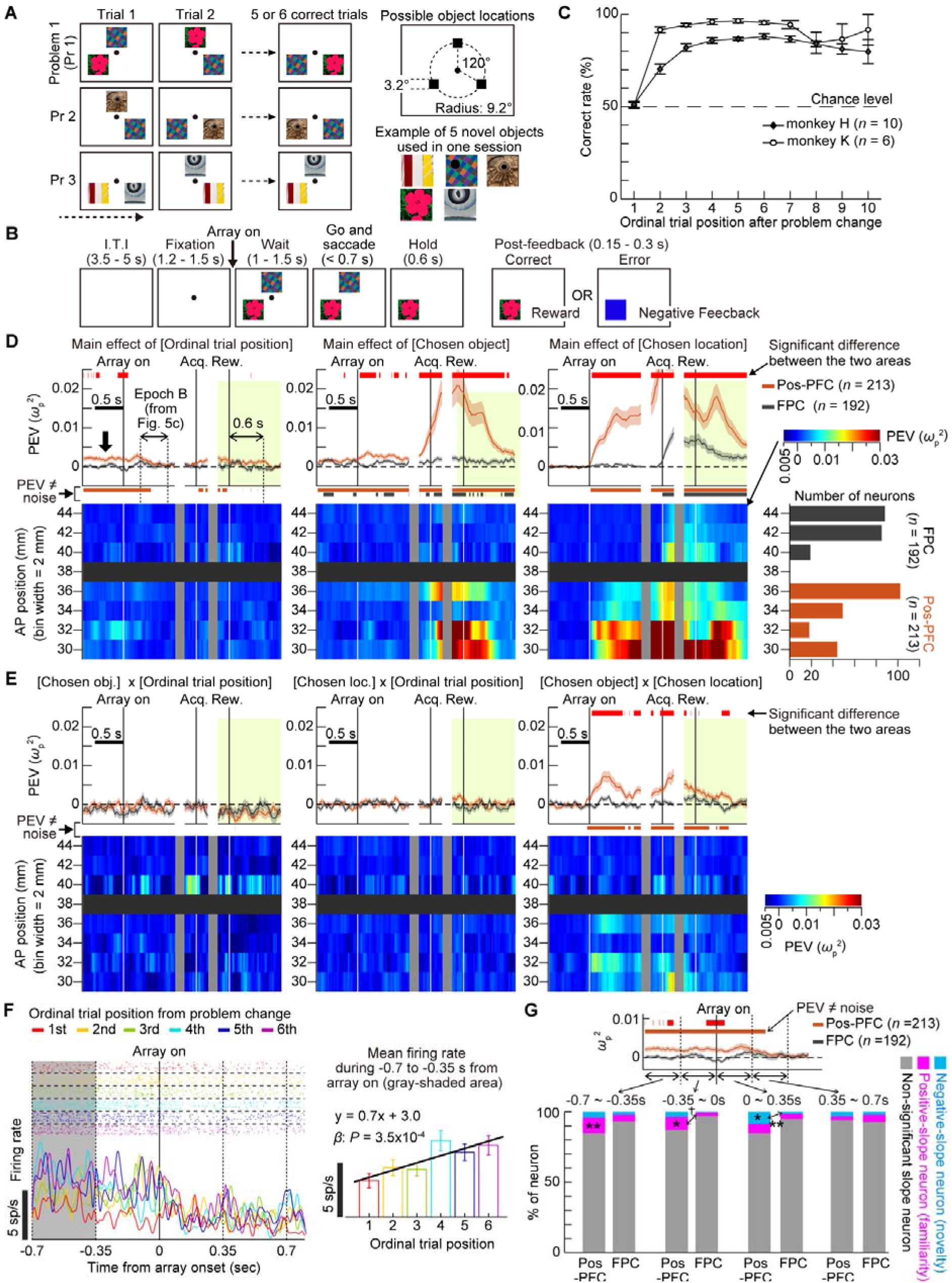
Neural activity related to learning across the anteroposterior LPFC. (**A**) Left: Task structure. Top right: Three possible object presentation locations. Bottom right: An example set of 5 novel objects used in one session. (**B**) Event sequence of the small-set FsL task. (**C**) Behavioral performance. Monkeys K and H completed an average of 78.7 and 69.6 problems per session. (**D**) Time course of population PEV (mean ± s.e.m.) for the three main effects in the pos-PFC (orange) and FPC (gray). For reference, the leftmost panel shows the duration of Epoch B as defined in Experiment 3 (Figure 5C, middle panel). Conventions as in Figure 5C. Bottom panels show PEV time courses at each 2-mm AP location. (**E**) Same as in **D**, but for the first-order interaction terms. Obj., object; loc., location. (**F**) Activity of a representative positive-slope neuron in the pos-PFC across trials 1 to 8 after problem change. Right panel plots the mean firing rate during the epoch from −0.7 to −0.35 s relative to array onset (grey-shaded area in left panel). (**G**) Percentage of negative-slope (cyan) and positive-slope neurons (magenta) in the pos-PFC and FPC across four analysis epochs: two before and two after array onset. Inset: An enlarged view of the PEV plot for the factor ordinal trial position shown in **D** (leftmost panel) near array onset.

Both monkeys performed significantly above chance (50%) from the second trial after problem change, indicating rapid learning of new S-R relations (**Figure 6C**, one-sample *t*-test). We calculated population PEV time series using a three-way ANOVA with factors: ordinal trial position from problem change (6 levels: 1st to 5th, and ≥6th), chosen object (5 levels), and chosen location (3 levels) (**Figures 6D** and **6E**). Correct and error trials were analyzed together, except during the peri-reward period (green-shaded area), where only the 1st to 6th *correct* trials after problem change were analyzed. In both FPC and pos-PFC, the PEV for the critical interaction term (chosen object × ordinal trial position) did not significantly increase relative to noise at any point (**Figure 6E**, leftmost panel, one-sample permutation test, FDR-corrected), indicating that selectivity for chosen object did not differ between exploration and exploitation phases. Thus, unlike the lateral orbitofrontal cortex (LOFC)^31^ and anterior cingulate cortex^32^, neither FPC nor pos-PFC contributed significantly to credit assignment, a key process in learning new S-R relationships.

Notably, significant PEV for the main effect of ordinal trial position appeared only in the pos-PFC (**Figure 6D**, leftmost panel), emerging well before array onset (arrow) and disappearing by 480 ms after. This contrasted markedly from Experiment 3, where significant PEV for this factor emerged only *after* array onset, in *both* the FPC and pos-PFC (**Figure 5C**, middle panel). Additional analyses revealed that, in the pos-PFC, this pre-array-onset ordinal-trial-position selectivity was primarily driven by an increase in positive-slope neurons signaling the occurrence of familiar contexts (**Figures 6F** and **6G**). Specifically, in the two pre-array analysis time windows (−0.7 to −0.35 s and −0.35 to 0 s; **Figure 6G**), the percentages of positive-slope neurons (magenta; 10.8% and 8.9%, respectively) significantly exceeded chance (2.5%) (*p* = 0.004 and *p* = 0.03, *p*-values corrected as in **Figure 5E**). However, in the 0–0.35 s window after array onset, the percentages of negative- and positive-slope neurons in the pos-PFC reversed (8.5% and 6.6%, respectively), both matching the result from Epoch B in Experiment 3 (**Figure 5E**; 7.2% and 8.7%, respectively; *p* > 0.39 for both across-experiment comparisons). These results indicate that, despite the large difference in object set sizes between Experiments 3 and 4, the pos-PFC flexibly adjusted its activity to distinguish exploration and exploitation. In contrast, the FPC showed no such flexibility, suggesting that it is not a critical node for learning in general. Another key observation was the significant difference in PEV for the factor chosen object between the two regions (**Figure 6D**, middle). Near reward time, the pos-PFC showed a robust increase in object selectivity, suggesting it closely monitored the chosen object’s identity. By contrast, the FPC weakly represented chosen object identity, only occasionally reaching significance and to a significantly lesser degree than the pos-PFC.

In addition, only the pos-PFC exhibited significant PEV for the interaction term (chosen object × chosen location) (**Figure 6E**, rightmost panel), representing ‘what in where’ information^33^ throughout the trial, with two prominent peaks after array onset and target acquisition. In contrast, even in a task where only object information was relevant, the FPC’s strongest representation near reward time focused only on chosen *location* (**Figure 6D**, rightmost panel). This suggests that, regardless of behavioral context, the monkey FPC minimally encodes object information. Since real-life learning cannot rely solely on spatial information, these results suggest that the monkey FPC is far from a critical hub for rapid learning, whether in novel contexts or involving reversals.

### Experiment 5: Testing FPC’s role in self-generated decision monitoring

Thus far, regardless of task demands, whether emphasizing spatial or object information, or involving functions previously implicated in the monkey FPC, the FPC has primarily and inflexibly engaged in retrospective coding of the chosen location associated with behavioral reports. This contrasts with the proposed role of the monkey FPC in decision monitoring, as suggested by Tsujimoto et al.^11^. In their spatial cued-strategy (CS) task, monkeys tracked a rewarded location (RL) from two possible targets (left or right) by using memory of the RL from the previous trial and a strategy cue indicating whether the RL would stay the same or shift. They termed this a ‘self-generated’ decision because it relied on internal memory of the RL. After such decisions (saccadic choice), they reported that FPC neurons exhibited significant chosen-location selectivity, sustained until both standard (0.5 s after target acquisition) and delayed (1.0 s after acquisition) reward timings. Notably, after externally-instructed action selection in the MGS (control condition), where RL memory carry-over across trials was unnecessary, significant chosen-location selectivity lasted only until the standard reward but diminished before the delayed reward, just 0.5 s later. Based on this duration difference, the authors interpreted the FPC’s peri-reward activity as reflecting a monitoring signal of self-generated decisions, integrating internal and external information to promote adaptive behavioral control, rather than merely encoding the chosen location of just-executed actions.

To resolve this discrepancy, we introduced a novel object-based CS task in which decisions were decoupled from space (**Figure 7A**). In this task, the monkey tracked the identity of a rewarded *object* (RO) and used a strategy cue to determine the RO on each trial (see **Methods** for task details). If FPC activity truly monitors ‘self-generated’ decisions and evaluates their outcome, its peri-reward activity should be selective to the chosen object identity rather than its location. For comparison, we employed the spatial CS task having the same temporal structure (**Figure 7C**) in which the monkeys tracked a rewarded location (RL) among two placeholders (RL and non-rewarded location, NL). The spatial and object CS tasks, along with the MGS task were performed in alternating blocks (counterbalanced across sessions). We recorded activity from 362 FPC and 400 pos-PFC neurons during all three tasks, and 72 FPC and 34 pos-PFC neurons only during the object CS and MGS tasks (recording locations, **Figure 1L**).

**Figure 7.**
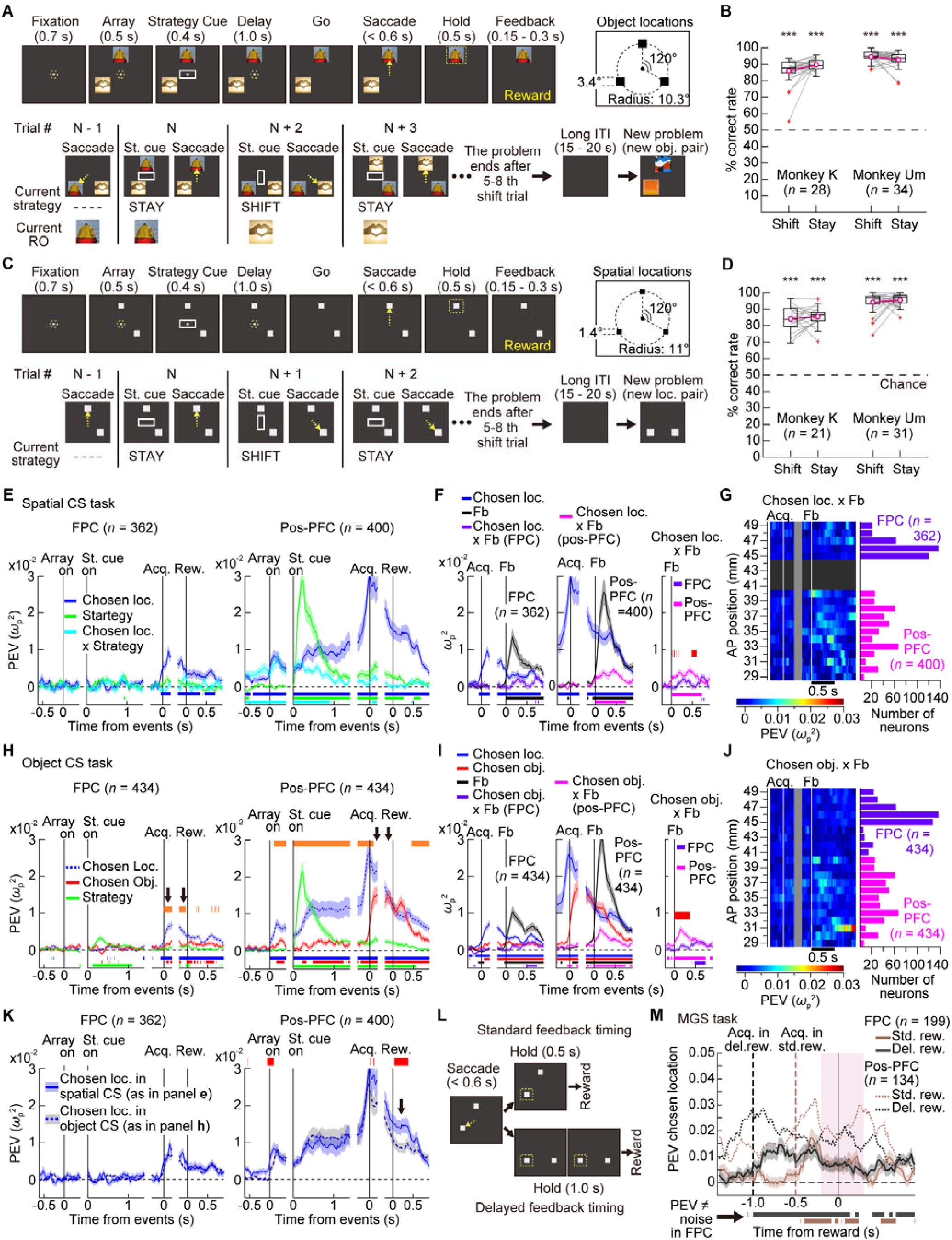
Neural activity related to spatial and object self-generated decision-making across the anteroposterior LPFC. (**A**) Object cued-strategy (CS) task. Top: Event sequence of a trial and three possible object locations. Bottom: Example task progression (correct trials only), showing the relationship between the strategy cue and the rewarded object (RO). The same object pair was avoided in successive problems. The first trial of each problem was always a stay trial with a randomly chosen RO. (**B**) Behavioral performance in the object CS task. (**C**, **D**) Same as in **A** and **B**, respectively, but for the spatial CS task. (**E**) Spatial CS task. Time course of population PEV (mean ± s.e.m.) for chosen location (blue), strategy (green), and interaction (cyan) in the FPC (left) and pos-PFC (right). Data are aligned to the onsets of array and strategy cue (st. cue), saccadic target acquisition (acq.) and reward (rew.). Lower colored horizontal bars mark periods of significant PEV in each factor. (**F**) Spatial CS task. Time course of population PEV for chosen location (blue), feedback (black), and their interaction in the FPC (left) and pos-PFC (middle). Data are aligned at acq. and feedback (fb). Lower colored horizontal bars mark periods of significant PEV in each factor. The rightmost panel compares the interaction PEV (chosen location × feedback) between the FPC (purple) and pos-PFC (magenta). Upper red horizontal bars mark significant differences between the two areas. (**G**) Spatial CS task. Time course of the interaction PEV (chosen location × feedback) across 1-mm AP locations. (**H**) Object CS task. Time course of population PEV for chosen location (dotted blue), chosen object (red), and strategy (green) in the FPC (left) and pos-PFC (right). Upper orange horizontal bars mark significant differences between PEV for chosen location and chosen object. Other conventions as in **E**. (**I**) Object CS task. Same as in **F**, but for PEV for chosen location (blue), chosen object (red), feedback (black) and the critical interaction term (chosen object × feedback). (**J**) Object CS task. Same as in **G**, but for the interaction PEV (chosen object × feedback). (**K**) Comparison of PEV for chosen location between the spatial and object CS tasks. Upper red horizontal bars mark significant differences between the two tasks. (**L**) Task progression after a correct saccade in the MGS and spatial CS tasks, in the standard and delayed reward conditions (blocked). (**M**) MGS task. Population PEV for chosen location (i.e., cue location), aligned at reward delivery in the FPC (solid lines) and pos-PFC (dashed lines). Lower horizontal bars mark time periods in which PEV in the FPC was significantly different from the noise level, for the standard (brown) and delayed (black) reward conditions. Pink-shaded area indicates peri-reward period (–0.2 to 0.3 s).

Both monkeys performed significantly above chance in both shift and stay trials of the object CS (**Figure 7B**, *p* < 7.6 × 10^−6^) and spatial CS tasks (**Figure 7D**, *p* < 1.2 × 10^−4^). At the neural level, we first examined population PEV in the spatial CS task using a two-way ANOVA (chosen location: 90°, 210°, 330°; strategy: stay/shift; **Figure 7E**). The results resembled prior studies^11,34^. In the FPC (left panel), the PEV for chosen location (blue) was significantly elevated only during the peri-reward period (after acq), with all other factors remained non-significant throughout the trial. Critically, in the object CS task (**Figure 7H**), a three-way ANOVA (chosen object × chosen location × strategy) showed that, during the peri-reward period, FPC activity (left panel) encoded the chosen location (blue) significantly more strongly than the chosen object (red) (arrows). Despite the task’s emphasis on object information, the chosen object PEV was weak and only sporadically significant, indicating that contrary to previous proposals, FPC peri-reward activity does not monitor the content of self-generated decisions but instead persistently encodes the chosen location retrospectively. In contrast, in the pos-PFC (**Figure 7H**, right), the chosen object PEV increased drastically, nearly matching that for chosen location, as evidenced by a non-significant difference between the two measures after acq (arrows).

For the 362 FPC and 400 pos-PFC neurons recorded during both CS tasks, only in the pos-PFC did the peri-reward PEV for chosen location significantly decrease in the object CS task compared to the spatial CS task (arrow, **Figure 7K**, right panel). This indicates that, in the object CS task, pos-PFC activity flexibly adapted to the change in task requirements, down-weighting the chosen-location signal and reallocating processing resources to encode the chosen object after acq. In contrast, FPC activity lacked such flexibility and consistently encoded chosen location with equal strength regardless of task-relevant modality. Additionally, unlike the pos-PFC (**Figure S6A**, right panel), the FPC (left) showed no significant PEV for interaction terms throughout the object CS trial, indicating little to no capacity for integrating object information during object-based self-generated decisions.

Next, to assess FPC’s role in decision outcome evaluation after feedback, we analyzed correct and error trials to determine if FPC activity discriminated RL vs. NL in the spatial CS task (**Figure 7F**, two-way ANOVA: chosen location × feedback) and RO vs. NO in the object CS task (**Figure 7I**, three-way ANOVA: chosen object × chosen location × feedback). For both critical interaction terms, [chosen location × feedback] in the spatial CS task and [chosen object × feedback] in the object CS task (rightmost panels), the FPC (purple) exhibited only a weak, delayed rise in PEV after feedback (fb), whereas the pos-PFC (magenta) showed an earlier and significantly stronger increase. A segmental analysis along the AP axis confirmed these results (**Figures 7G** and **7J**). Furthermore, in the object CS task, the three-way interaction PEV also rose earlier and significantly more strongly in the pos-PFC than in the FPC (**Figure S6B**). These findings indicate that the pos-PFC robustly and promptly evaluates decision outcomes across both spatial and object domains, while the FPC’s involvement is minimal and delayed, likely reflecting inputs from the pos-PFC or other regions.

Returning to Experiment 4 (**Figure 6**), we examined whether the same phenomena occurred during the feedback period in the small-set FsL task in which monkeys evaluated whether the chosen object was an S^+^ or not. Again, the PEV for the interaction term [chosen object × feedback] in the pos-PFC rose earlier and significantly more strongly than in the FPC whose increase was negligible (**Figure S7**). These results further emphasize the pos-PFC’s dominant role in decision evaluation across tasks and modalities, compared to the FPC.

Finally, in separate sessions (*n* = 23, monkey Um), following the prior study^11^ we introduced a delayed feedback condition (1.0 s after acq) for both the spatial CS and MGS tasks (**Figure 7L** and **8A**; 199 FPC and 134 pos-PFC neurons recorded in both tasks). To match the spatial CS task, the MGS task was slightly modified by adding placeholders (**Figure 8A**, bottom row). Contrary to the prior study’s conclusion that FPC’s prolonged (up to 1 s) chosen-location selectivity during the peri-reward period was exclusive to the spatial CS task, we found significant chosen-location selectivity in the FPC (and pos-PFC) continuing until the delayed reward (up to 1 s) in both MGS and spatial CS tasks (**Figure 7M**; for detailed results, including representative single-unit data, see **Figure 8**). Note that the prior study’s conclusion was based on 34 FPC neurons, used to assess whether the chosen-location selectivity persisted until the delayed reward in the MGS task, with just eight recorded in both tasks contributing to the time-course analysis (Fig. 7c,d in Ref. 11). Furthermore, the MGS task in Experiment 1 (**Figure 2**) also included both standard (250 ms) and delayed (600 ms) feedback conditions. Despite not involving self-generated decision-making, we again found in the MGS, that significant chosen-location selectivity in the FPC (and in the pos-PFC) persisted until the delayed reward (737 FPC and 736 pos-PFC neurons recorded in both reward conditions; **Figure S8**). These results suggest that simply tracking the duration of significant chosen-location selectivity after behavioral response is insufficient to assess FPC’s role in monitoring self-generated decisions. Additionally, the data from the single attention task in Experiment 1 (**Figure 2A**) which also included immediate and delayed (0.4 – 1.0 s) rewards, further support (in a manual response modality) that FPC’s peri-reward activity merely reflects spatial aspects of just-executed actions rather than serving as a monitoring/evaluation signal for integrative self-generated decisions (**Figure S9** and **Supplementary Text**).

**Figure 8.**
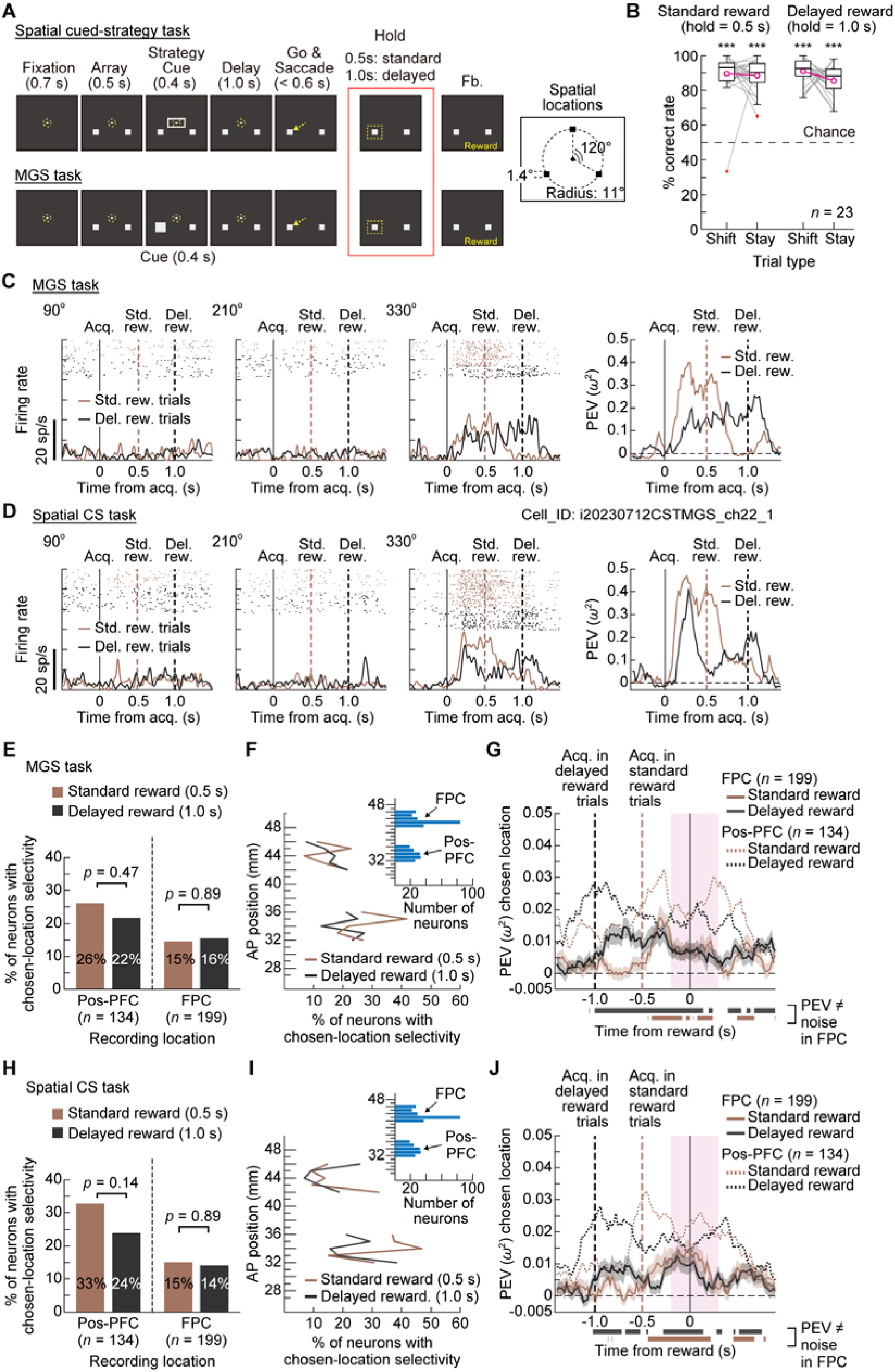
FPC neurons showed sustained chosen-location selectivity until delayed reward even in the MGS. (**A**) Standard (0.5 s hold period) and delayed (1.0 s) feedback conditions in the spatial CS and MGS tasks. In this version of the MGS task, two placeholders were introduced to match the display sequence in the spatial CS task. During the cue period, a memory cue (twice the placeholder size) appeared randomly on one of the two placeholders. MGS trials were grouped into a problem comprising 12-16 correct trials. Two placeholder locations stayed the same within each problem. (**B**) Behavioral performance of monkey Um in the spatial CS task under the standard (left) and delayed (right) feedback conditions. Performance in all four task conditions was significantly above chance (Wilcoxon signed-rank test; *p* < 0.0002, corrected). (**C**) MGS task: Raster-histograms in each cue location (three left panels) and the PEV for chosen location (cue location) aligned at saccadic target acquisition (acq.) for a representative FPC neuron. Data for the standard and delayed reward conditions are shown in brown and black, respectively. Brown and black vertical dashed lines indicate the onset of the standard (std.) and delayed reward (del. rew.), respectively. (**D**) Same neuron as in **C**, but for the spatial CS task. (**E**) MGS task: Percentage of neurons with significant chosen-location selectivity in the peri-reward period (−0.2 to 0.3 s from reward; pink shaded area in **G**) for the standard and delayed reward conditions in the pos-PFC (two left bars) and FPC (two right bars). *P*-values are for Fisher’s exact test (uncorrected). (**F**) MGS task: Percentage of neurons with significant chosen-location selectivity in the peri-reward period in each 1-mm segment along the AP axis, separately shown for the standard (brown) and delayed (black) reward conditions. No significant difference was found between the two conditions across all 10 segments (Fisher’s exact test, FDR-corrected). Inset shows the number of neurons recorded. (**G**) MGS task: Time course of population-averaged PEV for chosen location, aligned at reward delivery, in the FPC (solid lines) and pos-PFC (dotted lines). For clarity, the SEM (shaded area) is shown only for the FPC. Brown and black vertical dashed lines indicate timings of acq. for the standard and delayed reward conditions, respectively. Lower horizontal bars indicate time periods of significant PEV in the FPC for each condition (one-sample permutation test, FDR-corrected). The pink shaded area indicates the peri-reward period. (**H** to **J**) Same as in **E** to **G**, but for the spatial CS task. In panel **I**, across all 10 segments, no significant difference was observed between the two reward conditions.

## DISCUSSION

The monkey FPC has been commonly viewed as central to adaptive behavioral control, particularly in complex, non-routine environments. However, this notion has been shaped largely by limited experimental conditions, as well as the absence of direct comparisons with other prefrontal areas. Here, by intensively sampling neural activity throughout the anteroposterior LPFC, we re-defined the FPC’s role within the broader LPFC. Anteriorly, neural engagement progressively weakened, and was weakest in the FPC itself, for cognitive orchestration, rapid novel learning, and self-generated decision monitoring, functions that have previously been considered to be core to the leading role of the monkey FPC in adaptive control. FPC’s peri-feedback activity, previously considered to be crucial for information integration, was shown to be strikingly inflexible, persistently encoding predominantly the chosen location of a just-executed action regardless of the task demands. This rigidity was in stark contrast to the flexibility of the posterior-to-mid LPFC, the activity of which dynamically adjusted in response to multiple cognitive demands. Importantly, in novel learning and self-generated decision monitoring, we first replicated key observations from earlier FPC studies, validating our experimental approach. Nevertheless, by incorporating novel tasks and expanding LPFC sampling, we gained a more comprehensive perspective on the role of the FPC relative to the broader LPFC. Together, these findings reveal a reversed functional gradient in the monkey LPFC, characterized by posterior dominance, which challenges prevailing anterior-dominant models of LPFC function across primates. We begin by discussing each experiment individually.

In Experiments 1 and 2, dual-task-specific modulations of neuronal activity emerged only in the pos-PFC (**Figures 3E**-**3I** and **Figure 4I**). A sole prior study examining the role of the monkey FPC in cognitive orchestration^8^ inserted a secondary task (face-detection or free-reward consumption task) into ITIs of the Wisconsin card sorting task (WCST). Counterintuitively, FPC-lesioned monkeys outperformed controls in post-insertion WCST trials (though in both groups, post-insertion WCST performance was worse than that under no-insertion conditions). The authors proposed that the FPC is critical for disengaging from the ongoing task and reallocating cognitive resources to other potential goals in the environment. With an FPC lesion, monkeys were less concerned about extra-WCST events and stayed more engaged in the WCST during secondary-task insertion, thereby showing better performance in post-insertion WCST trials. However, several points warrant consideration before accepting this account. First, if the FPC truly reallocates resources to the interrupting task, FPC lesions should impair performance of that task. Yet, no quantitative comparisons were provided; it was only noted that both lesioned and control monkeys performed face detection at over 90% accuracy and consumed all the reward, implying little to no resource-reallocation deficit. Second, the FPC should also be needed to switch resources back to the WCST, and thus FPC-lesioned monkeys should struggle in returning to the WCST, making their superior post-insertion performance unlikely. However, this was not the case, and the absence of such reallocation deficits which suggests only a unidirectional resource allocation from the WCST to secondary tasks, remains unexplained.

In our study, if FPC activity was directing cognitive orchestration, it should at least distinguish between single- and dual-task conditions. However, in Experiments 1 and 2, it did not. Instead, neural activity reflecting dual-task-specific processes and adaptive resource allocation emerged mainly at AP levels corresponding to the posterior-to-middle thirds of the principal sulcus, declined anteriorly, and was absent in the anterior third and beyond (**Figures 3G** and **3I** and **Figure 4I**). Along with the lack of significant interaction effects in the FPC throughout Experiments 1–5, this suggests that the monkey FPC contributes minimally to multiple-goal management and information integration during cognitive orchestration. Rather, these processes are more strongly supported by the posterior-to-mid LPFC.

In novel learning (Experiment 3), 8% of FPC neurons responded preferentially to the appearance of novel objects (negative-slope neurons, **Figures 5D-5F**), mirroring the results reported by Nougaret et al.^13^ including the magnitude of percentage. At first glance, the FPC appears to be crucial for novel learning. However, we have demonstrated that in the pos-PFC, a significantly higher proportion of negative-slope neurons responded to novel objects at an earlier timing than in the FPC (**Figure 5E**). Furthermore, while reward-prediction-error (RPE) neurons which support novel S–R learning emerged in the FPC during the post-reward period, their prevalence was significantly greater in the pos-PFC (**Figure 5H**). In Experiment 4, when the object set-size was drastically reduced from that in Experiment 3, the pos-PFC adaptively changed activity patterns and remained engaged in learning, whereas the FPC showed no involvement (**Figures 6D** and **6G**). These findings suggest that the FPC plays a much more minor role in learning processes than the pos-PFC. Nevertheless, given that the ventral corticostriatal circuit, including OFC, is typically the primary driver of value-based object learning^23,31^, even the posterior LPFC’s role seems to remain relatively modest.

In the spatial and object CS tasks (Experiment 5), our findings challenge a prior proposal^11^ that the FPC is crucial for monitoring and evaluating self-generated decisions to guide future choices in dynamic environments. In our novel object CS task emphasizing object information, FPC activity near the feedback time, which was previously thought to reflect decision monitoring and outcome evaluation, failed to selectively encode chosen objects or their correctness (**Figures 7H-J**). Instead, it only represented task-irrelevant chosen-location information. In contrast, pos-PFC activity robustly encoded both monitoring and evaluation processes across spatial and object CS tasks, while also exhibiting flexible resource allocation by shifting the weighting of representations across modalities (space and object) in response to changing task demands (**Figure 7K**). Furthermore, contrary to earlier proposals^11^, an action need not be ‘self-generated’ to be represented in the FPC: comparable chosen-location-selective peri-reward activity appeared in the externally instructed MGS task and the spatial CS task under the delayed reward condition (**Figures 7M**, **8** and **S8**). A prior study on object coding in monkey FPC found minimal object selectivity during an object memory task^12^. Our findings confirm and extend this, showing that FPC neurons rarely exhibit object selectivity, even during initial phase of learning (Experiment 4) and self-generated decision-making (Experiment 5), contexts where the monkey FPC was previously considered crucial.

Our findings align with a line of anatomical evidence suggesting that the monkey FPC is not directly analogous to its human counterpart, a point often overlooked in discussions emphasizing cross-species similarities in prefrontal organization^3,35^. For example, (i) the human FPC is disproportionately expanded compared to those in apes and monkeys^36,37^; (ii) the monkey anterior LPFC lacks three major sulci present in the human lateral FPC^38^, suggesting the absence of a region analogous to the human lateral FPC, (iii) resting-state fMRI indicates that the functional connectivity of the human lateral FPC more closely resembles the mid-dlPFC than the FPC of the macaque^39^; and (iv) a tract-tracing study^40^ suggested that the monkey FPC corresponds predominantly to the medial portion of the human FPC which is subdivided into medial, orbital and lateral aspects. Furthermore, a meta-analysis of 148 tract-tracing studies places the monkey FPC relatively low in the prefrontal hierarchy, based on efferent-afferent and laminar-based connection asymmetries^41^. Our results that the monkey FPC (1) predominantly encodes action selection with highly limited object selectivity and minimal involvement in object-based decision-making, and (2) contains a moderate but significant proportion of reward-prediction-error neurons, suggest that the monkey FPC is an extension of the medial PFC which shares these characteristics^32,42,43,44^.

In summary, our findings challenge the notion that the monkey FPC is central to adaptive behavioral control in complex, non-routine environments, and instead highlight the posterior LPFC’s dominant role in functions previously attributed to the FPC. This raises broader questions regarding the conservation of prefrontal organization across primate species^45,46^. Specifically, we have demonstrated that the monkey LPFC exhibits a posterior-dominant functional gradient, reversing the anterior-dominant pattern widely assumed for humans. Consequently, evidence from the monkey FPC may be of limited relevance for understanding human prefrontal organization, particularly of its lateral-anterior aspect. The substantial anatomical and functional expansion of the human FPC, supporting uniquely human cognitive capacities such as mental simulation^47^, self-inference^1^, and hierarchical reasoning^27^, may represent an evolutionary addition beyond the explanatory scope of macaque models.

## METHODS

### Subjects

Experiments 1-5 were performed in four adult female macaques: two Japanese monkeys (*Macaca fuscata*, monkeys K and Um, weighing 8.4 (7 years old) and 7.5 kg (11 years old), respectively) and two rhesus monkeys (*Macaca mulatta*, monkeys T and H, weighing 4.6 (9 years old) and 6.0 kg (11 years old), respectively). Their ages are as of the start of the recording sessions. Different subsets of two to three monkeys participated in each experiment: Experiments 1 and 2 (K, T); Experiment 3 (K, H, Um); Experiment 4 (K, H); Experiment 5 (K, Um). All monkeys were housed individually under a 12-hour light/dark cycle (light: 8:30 a.m. to 8:30 p.m.). Prior to this study, monkey K had participated in two other posterior LPFC recording studies, which addressed research objectives distinctly different from those of the present investigation^27,54^.

### Ethics Statement

All experimental procedures were approved by the Animal Research Committee at the Graduate School of Frontier Biosciences, University of Osaka and complied fully with the guidelines of the National BioResource Project ‘Japanese Macaques’.

### Apparatus and Stimuli

During the experiments, the monkeys sat quietly in a primate chair within a dark, sound-attenuated shielded room. Their head movements were non-invasively restrained using a thermoplastic head cap made of standard splint material (MT-APU, 3.2 mm thick, CIVCO Radiotherapy, IA) ^25,48–50^. Visual stimuli were presented on a 17-inch TFT monitor positioned 50 cm from the monkey’s eyes. Eye movements were sampled at 120 Hz using an infrared eye-tracking system (ETL-200, ISCAN, MA). Fixation to the fixation point was controlled within a 6.5° square window (visual angle). Behavioral tasks were controlled by the TEMPO system (Reflective Computing, WA). The visual object stimuli used in Experiments 2–5 were obtained from the online gallery of the Art Institute of Chicago and the Open Images database^51^ and were cropped to the appropriate size.

### Implantation of recording chambers

Under general anesthesia and aseptic conditions, plastic recording chambers were stereotaxically implanted onto the lateral surface of the prefrontal cortex, guided by structural MRI (**Figure 1**). In monkey H (**Figure 1G**), a single cuboid chamber (internal dimensions: 12.0 mm width × 16.0 mm depth × 15.0 mm height; S-Company Ltd., Tokyo) was implanted, covering the frontopolar cortex (FPC) and extending posteriorly into the mid to posterior LPFC. For the remaining three monkeys, we implanted two chambers positioned along the anteroposterior axis: an anterior chamber over the FPC and a posterior chamber over the mid-to-posterior LPFC (pos-PFC, **Figure 1A**). The design of the FPC chamber varied among monkeys. For monkeys K (right hemisphere) and Um, a cuboid chamber was used (internal dimensions: 9.0 × 12.0 × 15.0 mm; S-Company Ltd). For monkey T, a cylindrical chamber was used (internal diameter: 8.0 mm; height: 18.0 mm). For the left hemisphere of monkey K, an elliptical cylindrical chamber was used (internal major/minor axes: 11.0/8.0 mm; height: 18.0 mm; O’hara & Co., Ltd., Tokyo). The design of the pos-PFC chamber also varied. Monkeys K (both hemispheres) and Um received cuboid chambers (internal dimensions: 12.0 × 16.0 × 15.0 mm; S-Company Ltd). For monkey T, we implanted a cylindrical chamber (internal diameter: 12.7 mm; product #: RC-T-S-P; Gray Matter Research, MT). In total, neural recordings were obtained from five hemispheres across four monkeys (**Figures 1C-1G**).

For surgical planning and identifying recording locations, we acquired high-resolution T1-weighted anatomical images in each monkey multiple times before and after chamber implantation under general anesthesia (medetomidine: 0.03 mg/kg, i.m.; midazolam: 0.2 mg/kg, i.m.; butorphanol tartrate: 0.3 mg/kg, i.m.). MRI data were obtained using a 3T Magnetom Prisma-Fit MRI scanner (Siemens, München, Germany) with a custom-built 12-channel phased-array receiver coil (Takashima Seisakusho Co., Ltd.). Preoperative anatomical scans typically used a 0.67 mm isotropic voxel size, while a 0.4 mm isotropic voxel size was used for identifying recording locations. Multiple anatomical scans were collected for each monkey as long as anesthesia remained effective, and the datasets were averaged to improve the signal-to-noise ratio. We reconstructed the monkey brains and recording sites using 3D Slicer software (version 5)^52^. Each monkey’s brain was semi-automatically rendered in 3D. Electrode tracks and their contact points on the brain surface were determined by aligning structural MRI images with the trajectory of the corresponding grid hole through which each track passed (**Figure 1B**). Coordinates for each contact point were determined using standard stereotaxic coordinates aligned to the orbitomeatal plane and expressed on the anteroposterior (AP) and mediolateral (ML) axes relative to the interaural midpoint (origin).

### Neural recording

We recorded all neurons encountered without any preselection. Across all monkeys and experiments, the FPC chamber yielded 140 unique sites in 155 total insertions (red and purple circles in **Figures 1C-1G**; AP coordinates ranged from 39.9 to 49.0 mm), while the pos-PFC chamber yielded 151 unique sites in 158 insertions (green circles; AP range, 26.5–40.8 mm).

For the pos-PFC recordings in monkey T (Experiment 1; **Figure 1F**, green dots), we used a unique setup consisting of a 32-channel semi-chronic microdrive system (SC-32, Gray Matter Research). This system housed 32 single-contact tungsten electrodes arranged in a grid pattern with an inter-electrode spacing of 1.5 mm. Approximately three hours before each session, monkey T was briefly transferred to the testing room, and each electrode was advanced individually by at least 65 μm (typically 130 μm) to ensure the isolation of new neurons.

For all other chambers, we used 24- or 32-channel acute linear multielectrode probes (U-Probe, Plexon, TX) with an inter-electrode spacing of 150 μm along a single shaft. Typically, a single U-Probe was inserted into each chamber using a custom-made grid designed for each chamber (φ0.6 mm holes, 1 mm grid spacing). To advance the U-Probes, we used an ultracompact micromanipulator (MO-903, Narishige, Tokyo) or a custom-built compact hydraulic microdrive (S-Company Ltd.). As in our previous studies^25,48^, we first punctured the dura using a guide tube (a shortened 24-gauge needle), then advanced the U-Probe into the cortex slowly in incremental steps, typically 500 μm at a time. The cortical surface was identified by monitoring pulsatile fluctuations (electrocardiogram) on superficial electrodes, and typically 3–5 superficial channels remained above the cortical surface during recordings. After insertion, we allowed 1–1.5 hours for recordings to stabilize, during which monkeys remained seated, watched nature and animal video clips, and received small snacks.

To ensure that differences in neuronal responses across recording locations were not due to systematic variations in signal processing or spike detection, we applied identical procedures to all recording chambers and monkeys. Raw extracellular neural signals were amplified and recorded using an RZ2 Bioamp Processor (Tucker-Davis Technologies, FL). Task-event information and eye-movement data were synchronized and transmitted to the RZ2. Neural signals were sampled at 24,414.08 Hz, and behavioral data were sampled at 1,017.25 Hz. Single-neuron activity was extracted by band-pass filtering the raw signals (300 Hz to 6 kHz), followed by semi-automatic offline spike sorting using Offline Sorter software (Plexon). During recording, action potentials were flagged for offline sorting if their waveforms’ trough or peak, typically the trough due to the extracellular recording method, crossed a threshold set at ±3.5 times the standard deviation (SD) of the raw signal’s continuously updated root-mean-square (RMS) value. This uniform cutoff method ensured a consistent signal-to-noise ratio (SNR) across all recording sites.

To confirm that the SNR was comparable across recording areas, we compared waveforms from all isolated single units recorded in the FPC and pos-PFC chambers using a standard method^53^. For each neuron, SNR was defined as the peak-to-trough amplitude of the mean waveform divided by the standard deviation of baseline noise, computed from the first 0.12 ms (i.e., the initial three data points before depolarization) of each waveform. Across Experiments 1-5, we isolated 2264 single units in the FPC and 2622 in the pos-PFC. Of these, 2260 units (99.8%) in the FPC and 2607 units (99.4%) in the pos-PFC met the criterion of SNR > 4, indicating excellent isolation quality in both recording locations.

### Statistical analysis

In this study, all population-level analyses included all recorded neurons without any prescreening procedure, except in the population histograms in **Figure 5J**, which included only neurons meeting specific criteria. All statistical tests were two-tailed and conducted in MATLAB (MathWorks). To address inflated Type I error rates in multiple hypothesis testing, we applied either the Benjamini-Hochberg (BH) procedure^54^ to control the false discovery rate (FDR) or the Holm’s sequentially rejective Bonferroni (Holm-Bonferroni) procedure, as appropriate. Corrected *p*-values are reported unless otherwise noted.

### Behavioral tasks and analysis

Details of behavioral tasks and analyses for each experiment are described primarily in the main text, with additional information provided below.

Experiment 1:

Monkey T completed 155 ± 34 SMT and 184 ± 29 DMT correct trials per session, whereas monkey K completed 143 ± 24 SMT and 176 ± 37 DMT correct trials (mean ± SD). In the attention task, a custom-made lever for behavioral responses was attached to the front of the chair at the monkey’s chest height. The lever remained in place during the dual-MGS-task (DMT) blocks but was removed during the single MGS (SMT) blocks, as it was unnecessary for SMT performance, and to avoid unintended associations. In DMT trials, because our primary focus was to examine how MGS-related activity differed between SMT and DMT conditions, we ensured clear temporal separation of MGS-related activity from attention-task-related activity by inserting the memory cue at least 0.7 s away from any attention task event (catch change or target change) and always before a target change (**Figure 2D**). Within these constraints, we randomly selected timings of memory cue presentation (1.0 to 4.1 s or 1.0 to 5.0 s from the attention cue offset in short trials and long trials, respectively, of the attention task). Despite this 0.7 s gap, extending the MGS analysis window beyond that period (e.g., to 1–2 s after memory cue onset) caused overlap with attention task events in some trials. In such trials, only the neural activity before that attention task event was used for analysis.

For the analysis of attention task performance, we assessed whether task difficulty differed among the three attention task conditions (Up, Down, and FR) by comparing the percent correct rates and lever-release response times (RT) in single attention task trials. Because we were interested in all pairwise comparisons among these conditions, we performed a series of three pairwise comparisons using a paired-permutation test with the Holm-Bonferroni procedure. There were three types of errors in the attention task: (i) fixation break (FB) error before the target change, (ii) premature lever release before the target change, and (iii) failure to initiate lever release within 0.6 s after the target change. Only the second and third types of errors were considered when calculating percent-correct rates.

To determine whether dual-task performance affected MGS task performance, we compared percent-correct rates across the SMT condition and the three DMT conditions. Session-by-session percent correct rates were calculated by dividing the number of correct trials by the total number of trials in which the monkeys successfully reached the go-signal for the memory-guided saccade (i.e., the end of the follow-up fixation period, **Figure 2D**). A correct trial was defined as one that satisfied both of the following criteria: (i) successful saccadic target acquisition within 0.5 s after the go-signal, and (ii) successful gaze-holding at the correct placeholder for 0.25 or 0.6 s. Any other eye movement after the go-signal was considered an error (i.e., failure to acquire the correct target or failure to maintain fixation after acquisition). Since the SMT condition did not involve a target change or a subsequent lever release, these events were scheduled but executed as ‘empty events’ with no change in the physical stimuli or a behavioral response.

For the neuronal analysis in **Figures 3G** and **3I**, which shows the percentages of location-selectivity_dual>single_ and firing-rate_dual>single_ neurons, we used the following time windows for each task period: fixation (−0.6 to 0 s from the end of the fixation period), memory cue (0.1 to 0.4 s from cue onset), delay (0.4 to 1.0 s from cue onset), and pre-go (−0.3 to 0 s from the saccade go signal). Neural recording sites in Experiment 1 covered roughly 20 mm along the AP axis, encompassing the entire principal sulcal area and extending beyond it (from AP 26.5 to 46.1 mm in stereotaxic coordinates). Along the AP axis, 47 FPC sites (red markers, **Figure 1H**) spanned a 6.2 mm region (AP 39.9 to 46.1 mm), and 69 mid-to-posterior LPFC (termed pos-PFC) sites (green markers, **Figure 1H**) spanned 13.5 mm (AP 26.5 to 40.0 mm).

Experiment 2:

During the few-shot learning (FsL) task, a secondary task was inserted at a random point between the second trial and the seventh correct trial within a problem. Secondary-task insertions occurred twice per problem. On each insertion, one secondary-task condition was selected randomly. Each insertion was separated by at least two to three correct trials. Because the no-insertion condition involved no secondary-task insertion, 11% (1/9) of problems had no actual secondary-task insertion, 44% (4/9) of problems had one, and 44% (4/9) had two. Thus, the present design ensured enough trials for both the MGS and time-out/FR conditions. In the MGS task, there were six possible cue locations (30°, 90°, 150°, 210°, 270°, or 330°) on an imaginary circle with a 12° radius. To calculate FsL task performance (**Figure 4E**), we included all trials regardless of whether secondary-task insertions occurred before or after that trial, except those that were aborted due to fixation breaks prior to the go signal. For the three-way ANOVA of the FsL task (**Figure 4H**), we included ‘trial location’ as a factor. For its two levels, pre- and post-insertion, we averaged activity across two correct trials, before (−2, −1) and after (+1, +2) insertion, respectively. For the two-way ANOVA analyzing MGS activity (**Figure 4I**), 26 FPC neurons were excluded from the analysis due to a code error in the first session in monkey T that resulted in the cue location of the dual-task MGS being unrecorded.

Experiment 3:

In the modified FsL task, the location and size of object presentation were identical to those in Experiment 2 (see **Figure 4B**). In **Figure 5J** (normalized population histograms of negative-slope neurons), each neuron’s firing rate in successive 50-ms bins was normalized to its mean firing rate during the period from −1.0 to 1.0 s relative to the reward averaged across all trials. Because the statistical comparisons between different reward conditions were based on a within-subject design (i.e., paired permutation test), within-subject SEM^55,56^ was used for the error bars. Note that the comparison between the 1st reward (cyan) and the 8th reward (brown) was inherently expected to show a significant difference (1st > 8th), because the selection criterion for negative-slope neurons required a decrease in activity from the 1st to the 8th reward. Consequently, performing this comparison on neurons that meet this criterion constitutes a case of double-dipping. The remaining three comparisons, however, were valid because they were independent of this selection criterion.

Experiment 4:

Apart from the small-set FsL task, we recorded neural activity during the MGS task in separate blocks. The MGS task had the same parameters as in Experiments 2 and 3, except that there were only three possible cue locations (90°, 210°, and 330°) instead of six, to match the object presentation locations in the small-set FsL task. In the neural analysis of the small-set FsL task, the three-way ANOVA ([Ordinal trial position] × [chosen object] × [chosen location], **Figures 6D** and **6E**) used a reduced model (i.e., main effects and two-way interactions only) rather than the full model, owing to an insufficient number of trials to reliably estimate the three −way interaction.

Experiment 5:

In the object CS task (**Figure 7A**), each trial began with a 0.7-s fixation period, followed by the appearance of two objects at two of the three predetermined locations (array period). A strategy cue then appeared around the FP (strategy cue period). A horizontally elongated cue (ellipse for monkey K, rectangle for monkey Um; subtending 2.3°× 5.7°) indicated that the previously rewarded object (RO) remained rewarded (stay trial). A vertically elongated cue (rectangle with a flicker for monkey K, plain rectangle for monkey Um) indicated that the previously non-rewarded object (NO) became rewarded (RO), and the previous RO became NO (shift trial). After a 1.0-s delay period, the FP disappeared (go-signal), prompting the monkey to saccade to one of the two objects within 0.6 s and hold it for 0.5 s (hold period). The unchosen object disappeared upon selection. If the chosen object was the RO, the monkey received a reward. Otherwise, the chosen object turned blue (error signal), and correction trials with the same strategy cue continued until the monkey chose correctly. To ensure that the new RO was learned after switching, a correct shift trial was always followed by a stay trial; otherwise, shift/stay cues were randomized. Trials were grouped into problems, each ending once 5–8 shift trials were completed with the same object pair. A long ITI (15–20 s) then preceded a new problem with a new object pair (**Figure 7A**, bottom right). Each daily session introduced only five novel objects. Across 62 sessions, excluding correction trials, the two monkeys performed an average of 347.3 ± 79.2 trials per session (mean ± SD, including both correct and error trials), divided into 118.8 shift trials (34.2%) and 228.5 stay trials (65.8%).

The spatial CS task (**Figure 7C**) had the same temporal structure as the object CS task, but the monkeys tracked a rewarded location (RL) among two placeholders (RL and non-rewarded location, NL). Throughout each problem (consisting of 5–8 shift trials), the same location pair was used, selected from the 90°, 210°, and 330° locations. In each trial, choosing the RL earned a reward and choosing the NL turned that placeholder red (error signal). A correct shift trial was always followed by a stay trial. Over 52 sessions, excluding correction trial, the two monkeys performed 201.0 ± 62.5 (correct and error) trials per session (69.1 shift trials (34.4%) and 131.9 stay trials (65.6%)).

For behavioral analysis of the object and spatial CS tasks, we excluded (i) the first trial of each problem and (ii) correction trials. For neural analysis, we excluded (i), (ii), and error trials (unless otherwise noted). Thus, all trials included in the neural analysis were immediately preceded by a correctly performed trial. In the object CS task, the monkeys’ performance was as follows: monkey K, shift trial, 86.0 ± 8.0%, stay trial, 89.7 ± 3.4%; monkey Um, shift trial, 94.6 ± 3.3%, stay trial, 92.8 ± 4.5% (mean ± SD; chance = 50%) (**Figure 7B**). In the spatial CS task, their performance was as follows: monkey K, shift trial, 84.2 ± 7.8%, stay trial, 85.4 ± 5.7%; monkey Um, shift trial, 94.3 ± 6.2%, stay trial, 95.7 ± 3.8% (**Figure 7D**).

During the main sessions, the MGS task was similar to that used in Experiment 4, featuring three possible cue locations (90°, 210°, and 330°). However, the temporal progression differed: fixation period = 1.2 s, memory cue period = 0.25 s, delay period = 1.15 s, response period < 0.6 s, and hold period = 0.5 s. The interval from memory-cue onset to the go signal was 1.4 s, matching the duration from strategy-cue onset to the go signal in the CS tasks.

### Analysis of neural selectivity for task variables

To quantify the strength of neural selectivity for a given task variable, we used the proportion of explained variance (PEV). For our main analyses (**Figures 3**-**7**), we computed the partial ω² value for each factor in an *n*-way ANOVA model. The calculation was performed using the formula:

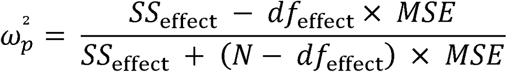

where *SS*_effect_ and *df*_effect_ are the type sum of square and the degree of freedom (between groups), respectively, for the factor of interest. *MSE* is the mean square error within groups, and *N* is the total number of trials. The partial ω² indicates how much variance in a neuron’s trial-by-trial firing rate is explained by a given factor, while accounting for the influence of other factors in the ANOVA model. For the analysis in **Figure S3, 8, S8 and S9**, with a single analytical variable, we computed the standard omega-squared (ω^2^) for a one-way ANOVA model using the following formula:

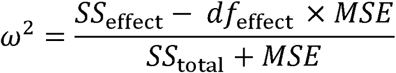

where *SS*_total_ is the total sum of squares for the entire data set. For the time-series analysis, PEV was computed using 200-ms time windows slid in 20-ms increments across the entire trial. In each time window, the observed population-averaged PEV values were compared against a null distribution created by randomly shuffling condition labels 10,000 times, thereby simulating the distribution of PEV values expected by chance (one-sample permutation test). The resultant permutation *p*-values were then corrected for multiple comparisons across time windows by controlling FDR under a BH procedure. The significance level was set at *p* < 0.05. Depending on the nature of the comparison (within-subject vs. between-subject), we additionally used paired or two-sample permutation tests as appropriate.

## ACKNOWLEDGEMENTS

The authors thank S. Kitazawa, J. Duncan and S. Funahashi for their comments on the manuscript. This work was supported by the Grants-in-Aid for Scientific Research (KAKENHI 18K03197 and 24K00506) from the Japan Society for the Promotion of Science (JSPS) to K.W., and by the ImPACT Program of the Cabinet Office, Government of Japan, to T.S. Japanese macaques were supplied by the National BioResource Project “Japanese Monkeys,” funded by the Ministry of Education, Culture, Sports, Science and Technology (MEXT), Japan.

## Data and materials availability

All raw spiking data and behavioral event data supporting the current study will be made publicly available upon peer-reviewed publication. Each neuron’s dataset includes its AP/ML recording coordinates, trial label, and fully timestamped behavioral events. Code for the behavioral and neuronal analyses will be made publicly available upon peer-reviewed publication. Any additional information required to reanalyze the result reported in this paper is available from the lead contact upon request.

## AUTHOR CONTRIBUTIONS

K.W. and T.S. conceived of the study. All authors designed the experiments. K.W. and M.H. performed the surgery. K.W. conducted experiments, analyzed the data, and wrote the initial draft. All authors discussed the results and reviewed the manuscript at all stages.

## DECLARATION OF INTERESTS

The authors declare that they have no competing interests.

## Supplementary Materials

### Supplementary text for Figure S2

One potential issue in the analysis of **Figures 3E, 3H** and **3I** (Experiment 1) is that in most recording sessions (60/75), the SMT and DMT conditions were conducted in relatively long, separate blocks. This setup may have introduced small but gradual changes in firing rate (drift) between the two conditions, potentially affecting the results, particularly the factor task that compared overall firing-rate differences between the SMT and DMT. To address this concern, we repeated the analysis using only neurons recorded during the frequent task-switching sessions (*n* = 15 sessions) in which SMT and DMT blocks alternated frequently (8 ± 1.2 times per session; every 40–50 correct trials). The results (**Figure S2**) closely matched those in **Figure 3**. The population PEV time courses (**Figure S2A)** were comparable to those in **Figure 3E**, and the time course of percentages of dual-task preferring neurons (**Figures S2B and S2D**) were also comparable to those in **Figures 3F** and **3H**. Notably, the percentages of firing-rate_dual>single_ neurons again showed a significant anterior decline (**Figure S2E**), as seen in **Figure 3I**. The percentages of location-selectivity_dual>single_ neurons also showed anterior decline, though not significant (**Figure S2C**). Additionally, as in **Figure 3H**, pos-PFC activity exhibited heightened readiness during the initial fixation period in DMT compared to SMT (arrows in **Figure S2D**), which likely reflected the expectation of higher mental workloads in DMT. These findings indicate that potential neuronal drift had negligible impact on our results.

### Supplementary text for Figure S5

One concern in the analysis of **Figures 5C-5F** (Experiment 3) is the inclusion of data from both fixation (Experiment 3) and non-fixation (Experiment 2) conditions before and after array onset in the FsL task. This could confound the neuronal results, as eye movement frequency after array onset may vary depending on the trial’s ordinal position following the problem change. Notably, this issue also applies to the previous FPC study, where fixation was not required at all throughout the Object-in-Place task^13^.

To address this concern, we repeated the analyses using only Experiment 3 data (**Figure 5A**, modified FsL task). The results (**Figure S5**) were essentially unchanged, ruling out eye movement frequency as a confounding factor. Specifically, for the factor ordinal trial position, the significant PEV elevation in the pos-PFC again preceded that in the FPC by 180 ms (vertical arrow, middle panel, **Figure S5A**). The percentages of negative- and positive-slope neurons during these slightly shifted Epochs A and B (100–280 ms and 280–760 ms from array onset, respectively) were comparable to those in **Figure 5E** (**Figure S5B**). Other findings (**Figure S5C**) also aligned with **Figure 5F**. These results indicated that the findings in **Figures 5C-5F** were not affected by factors related to eye movement.

### Supplementary text for Figure S9

To further test whether the FPC’s peri-reward activity truly monitors decision-making, we analyzed neural activity in the single attention task trials (Experiment 1) which also included two reward timing conditions: immediate and delayed (0.4 – 1.0 s from lever release) reward (**Figure S9A**). In the attention task, a single action (lever release) was used to report the outcome of a yes/no decision (as in the framework of the signal detection theory), judging whether a color change occurred at the ring prespecified by the attention cue. Thus, if FPC activity indeed monitors decision-making as previously suggested^11^, during the peri-reward period, it should distinguish between the three possible locations for target change, a critical decision variable, despite the decision being reported through a single action. All recorded neurons from Experiment 1, which included data from both reward conditions, were used in this analysis (FPC, *n* = 853; pos-PFC, *n* = 1219).

The time course of PEV for the target change locations, aligned to reward time (time 0 marks reward delivery; **Figure S9B**), indicated that FPC neurons (solid traces) did not discriminate the location of target change before or after the reward in either the immediate (pink) or delayed (green) reward conditions (FDR-corrected *p* > 0.36 for all 61 time bins in each reward condition). In contrast, pos-PFC neurons significantly discriminated the location of target change from before to until well after the reward in both the immediate and delayed reward conditions (dashed pink and dashed green trace, respectively).

When aligned to the timing of lever release (**Figure S9C**), the PEV in the pos-PFC exhibited a common peak before lever release across the two reward conditions, which likely reflected a location-selective visual response to the target change. After the reward, in the delayed reward condition (dashed green trace), the PEV in the pos-PFC remained significant well after the closest (earliest) possible reward timing (0.4 s after lever release, vertical dashed green line), with a much longer period of significance compared to the immediate reward condition (dashed pink trace) (see arrow). This late PEV component, the decay rate of which varied markedly between the two reward timing conditions, likely reflects retrospective monitoring of target change location, a critical decision variable in this yes/no decision-making. In contrast, the PEV in the FPC (solid traces) did not show significant values either before or after the reward in both reward conditions. The lack of location selectivity in FPC peri-reward activity is likely due to the use of a single action (lever release) for behavioral reporting, although the report itself was based on a location-based decision. This suggests that the location-selective peri-reward activity in the FPC, previously interpreted as a monitoring signal for location-based decisions, merely reflected the chosen saccade location used to express those decisions in that prior study^11^. These results further confirm that, while pos-PFC peri-reward activity retrospectively monitored the decision, FPC activity did not. Instead, FPC activity was limited to representing the spatial information of just-executed actions.

**Figure S1.**
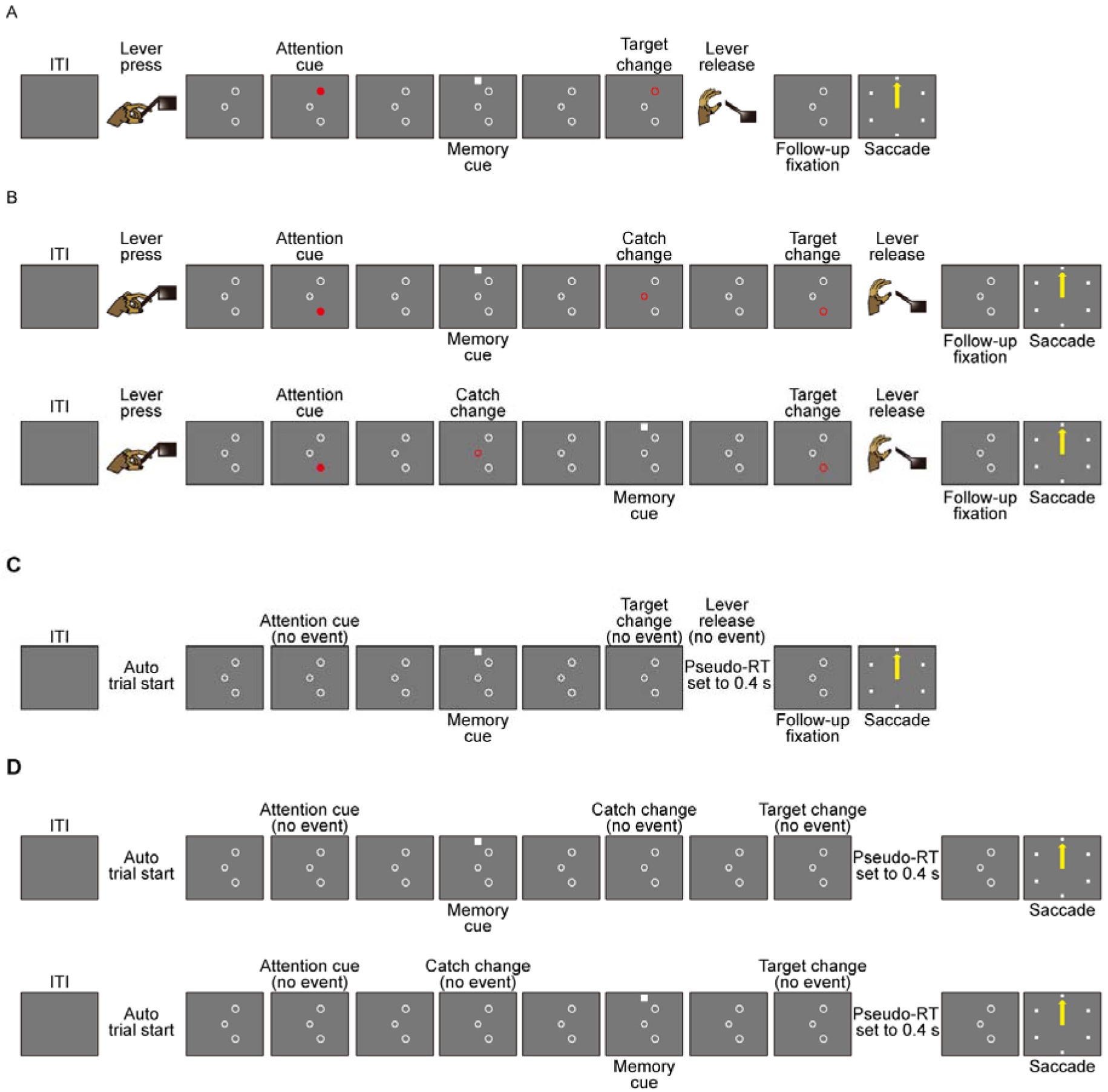
Example DMT and SMT trial sequences. (**A**) MGS task inserted into a short trial of the attention task. The MGS insertion occurred in 66.7% of all attention-task trials, with this type of DMT trial comprising half of them (33.3% of all attention-task trials). (**B**) MGS task inserted into a long trial of the attention task. The trial in the upper row shows MGS insertion during the wait1 period (before catch change), while the bottom row shows insertion during the wait2 period (after catch change). These DMT trials made up half of all DMT trials, accounting for 33.3% of all attention-task trials. (**C**) SMT trial matching the time course of the short attention task trial (as in **A**). In the SMT block, a memory cue was presented in every trial. This type of SMT trial comprised 50% of all SMT trials. (**D**) SMT trial matching the time course of the long attention task trial (as in **B**). This type of SMT trial also made up 50% of all SMT trials. In the SMT block, while all attention-task events were scheduled, they were executed as ‘empty events’ with no physical changes in stimulus. Trials were automatically initiated by the appearance of the fixation ring (FR) after an intertrial interval.

**Figure S2.**
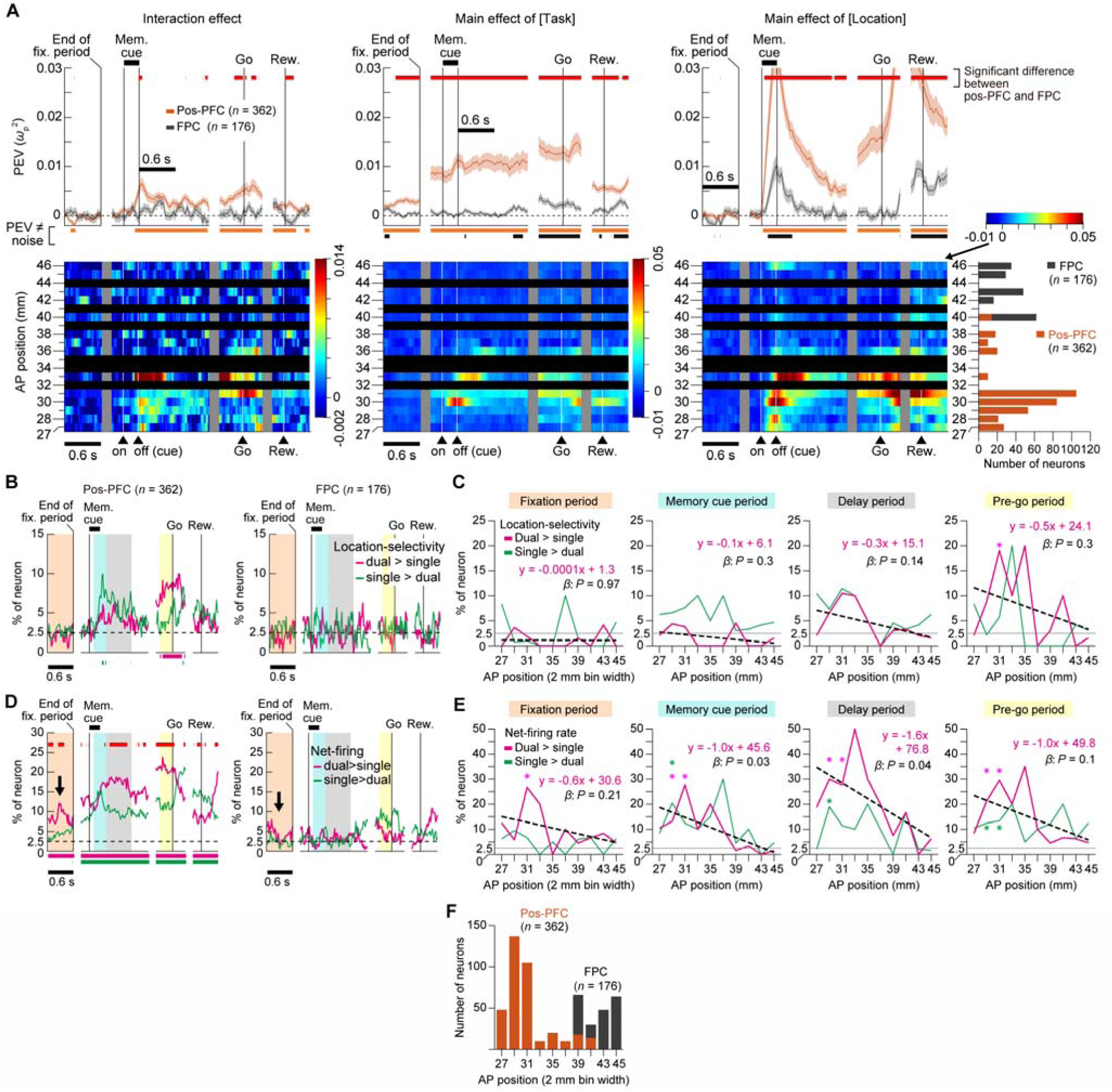
Neural activity during the frequent task-switching sessions in Experiment 1. (**A**) Upper row: Time course of population-averaged PEV for interaction (left), and main effects of task (middle) and location (right). For the pos-PFC analysis, data from the dorsal and ventral pos-PFC are combined. Bottom row: Time course of population-averaged PEV in each 1-mm segment of recording locations along the AP axis. Horizontal black strips in the plot area indicate locations where no neurons were recorded. Other conventions as in Figure 3E. (**B**) Time course of the percentages of the location-selectivity_dual>single_ and location-selectivity_single>dual_ neurons (magenta and green curves, respectively). Conventions as in Figure 3F. (**C**) Percentages of the location-selectivity_dual>single_ (magenta) and location-selectivity_single>dual_ (green) neurons in each 2-mm segment along the AP axis during the fixation (orange), memory cue (cyan), delay (gray) and pre-go (yellow) periods. Due to the small number of neurons, percentages were calculated using 2-mm bins. (**D** and **E**) Same as in **B** and **C**, respectively, but for the firing-rate_dual>single_ (magenta) and firing-rate_single>dual_ neurons (green). (**F**) Number of recorded neurons in each 2-mm segment along the AP axis.

**Figure S3.**
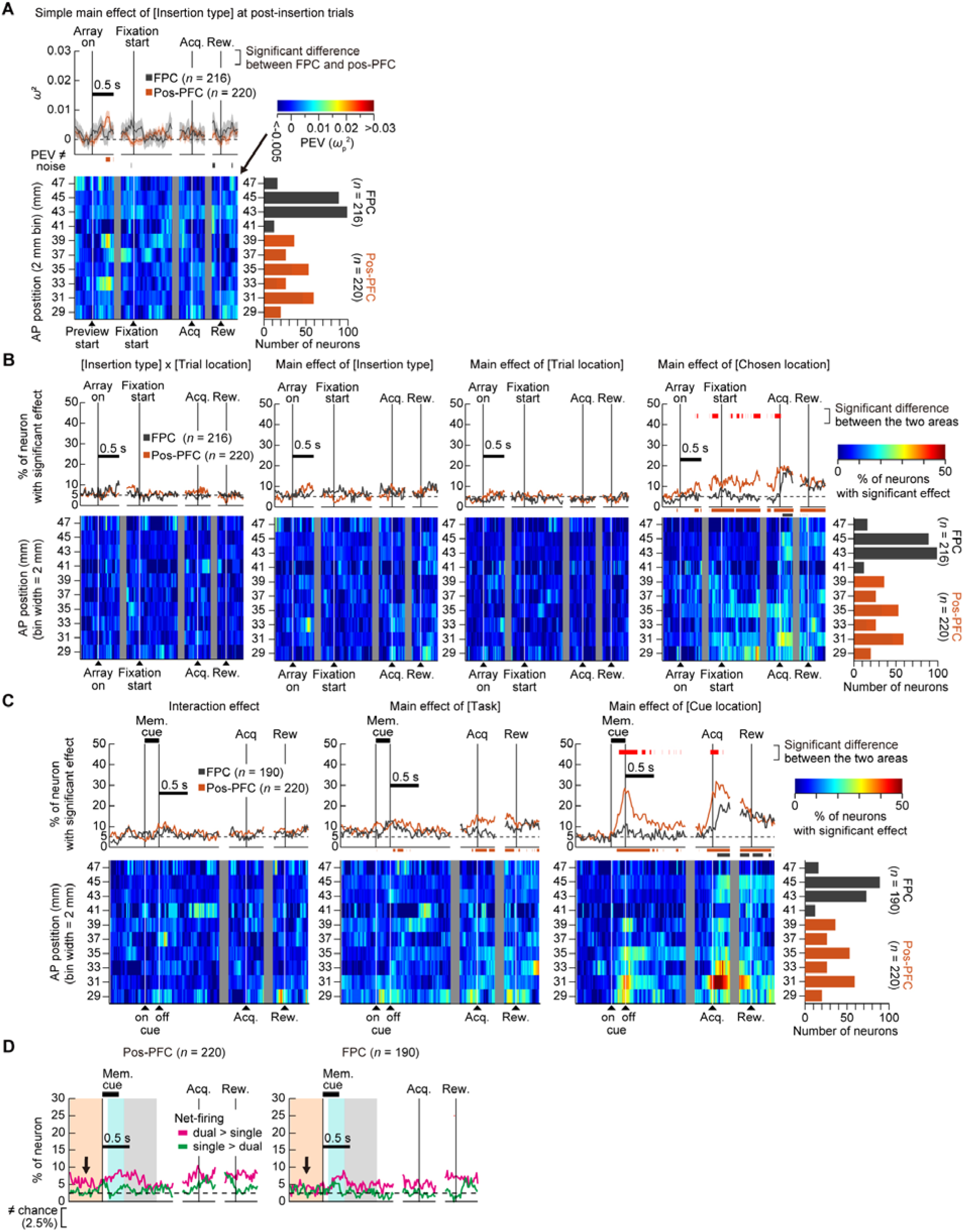
Neural activity during the serial multimodal dual task in Experiment 2. (**A**) Time course of population PEV for the simple main effect of the factor insertion type, using only post-insertion trials. Conventions as in Figure 4H. (**B**) Time course of the percentages of significant neurons for the four critical ANOVA terms in the FsL task, aligned at the start of the preview period (array on) and wait period (fixation start), and the timing of target acquisition (acq.) and reward (rew.). Top row: Red horizontal bars mark time periods with significant differences in the percentages between FPC (gray curve) and pos-PFC (brown curve) (Fisher’s exact test, FDR-corrected *p* < 0.05). Lower colored horizontal bars mark time periods in which the percentage was significantly different from chance (5%) in each recording area (Fisher’s exact test, FDR-corrected *p* < 0.05). Bottom row: Time course of the percentage of significant neurons in each 2-mm segment of recording locations. The rightmost panel indicates the number of recorded neurons. (**C**) Same as in **B**, but for the comparison between single-task and dual-task MGS. (**D**) MGS task: Time course of the percentages of the firing-rate_dual>single_ (magenta) and firing-rate_singlel>dual_ neurons (green) in the pos-PFC (left) and FPC (right, shown for reference). The result resembled the pattern observed in Experiment 1. In the pos-PFC, firing-rate_dual>single_ neurons consistently outnumbered firing-rate_single>dual_ neurons throughout the trial, though not significantly. Notably, during the fixation period, the pos-PFC showed an elevated (though not significant) percentage of firing-rate_dual>single_ neurons compared to firing-rate_single>dual_ neurons (arrows), suggesting that the pos-PFC activity was primed from the foreperiod for the anticipated higher cognitive demand of dual-task conditions.

**Figure S4.**
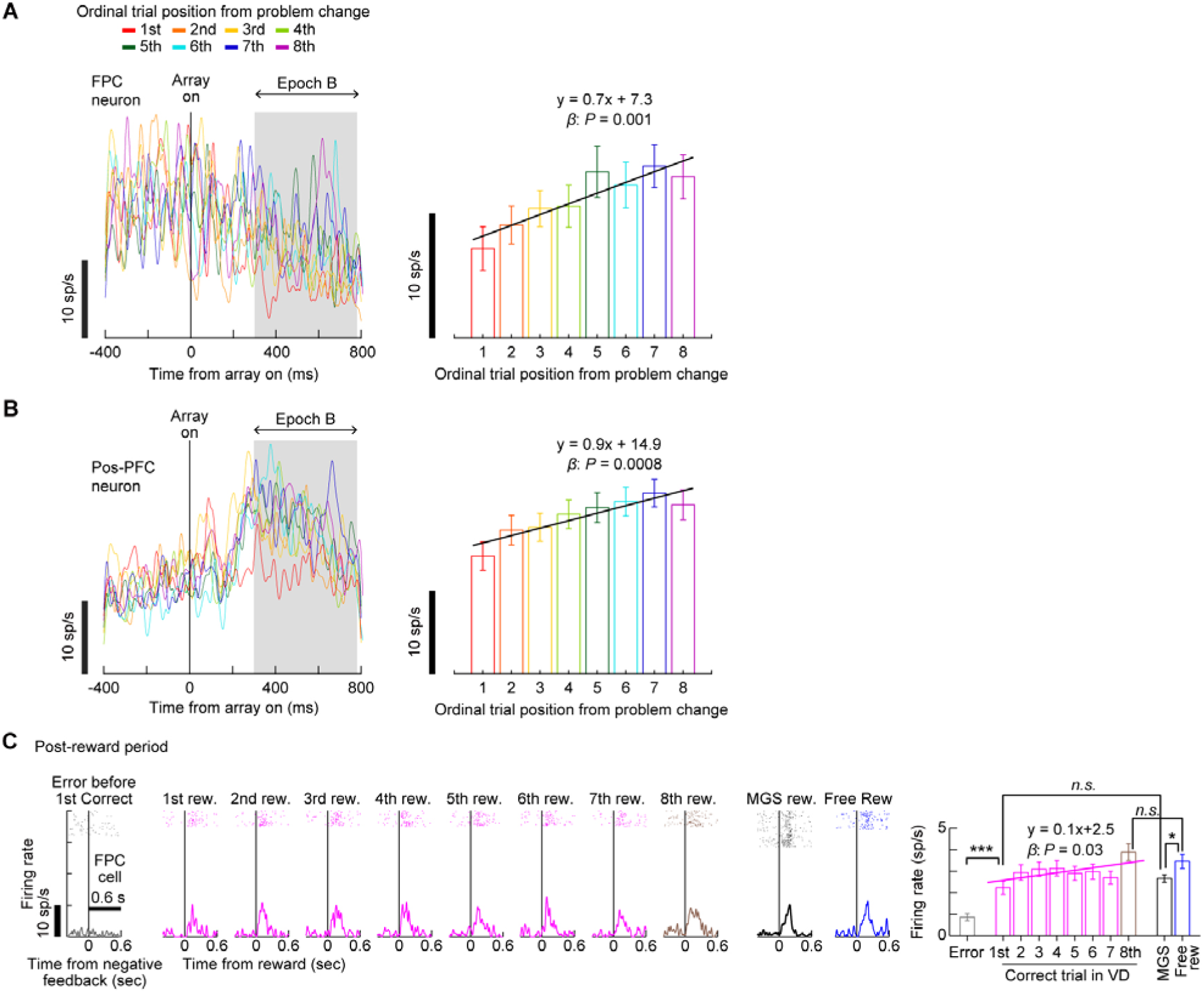
Neural activity during the serial multimodal dual task in Experiment 2. (**A** and **B**) Activity of representative positive-slope neurons after array onset in the FPC (**A**) and pos-PFC (**B**) across trials 1 to 8 after problem change. The right panel shows the mean firing rate in Epoch B (shaded area) with OLS regression results. (**C**) Post-reward activity of a representative positive-slope neuron in the FPC across 1st to 8th correct trials after the problem change, shown alongside activity in error trials (leftmost panel) and in MGS reward and free reward. The rightmost panel indicates the mean firing rate for each trial type in Epoch C (0 – 0.6 s from reward) with OLS regression results.

**Figure S5.**
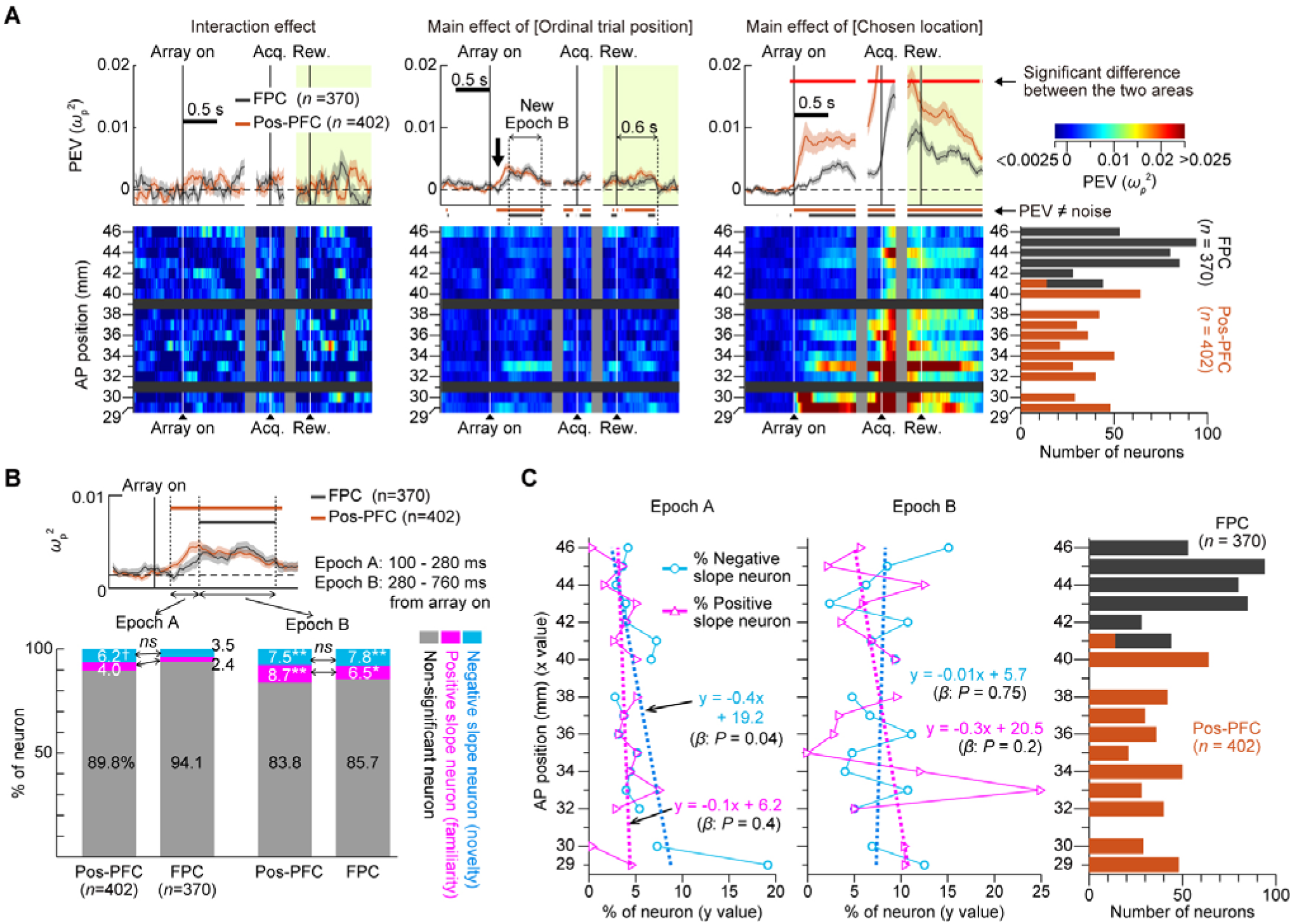
Neural activity across the anteroposterior LPFC in the modified FsL task (only data from Experiment 3 with fixation requirement before and after array onset). (**A**) Time course of population-averaged PEV. Conventions as in Figure 5C. (**B**) Comparison of the percentage of negative- (cyan) and positive-slope (magenta) neurons between the pos-PFC and FPC in Epochs A and B. Conventions as in Figure 5E. The inset PEV plot enlarges the region near ‘array on’ in **A** (second panel from left). (**C**) Percentage of negative- (cyan) and positive-slope(magenta) neurons across 1-mm segments along the AP axis for Epochs A (left panel) and B (middle panel). No FPC subregion exceeded the 2.5% chance level for either negative- or positive-slope neurons in either epoch (for both epochs: *p* > 0.28, *p*-values corrected for 16 comparisons in each epoch). Conventions as in Figure 5F. Rightmost panel shows the number of neurons recorded per 1-mm segment.

**Figure S6.**
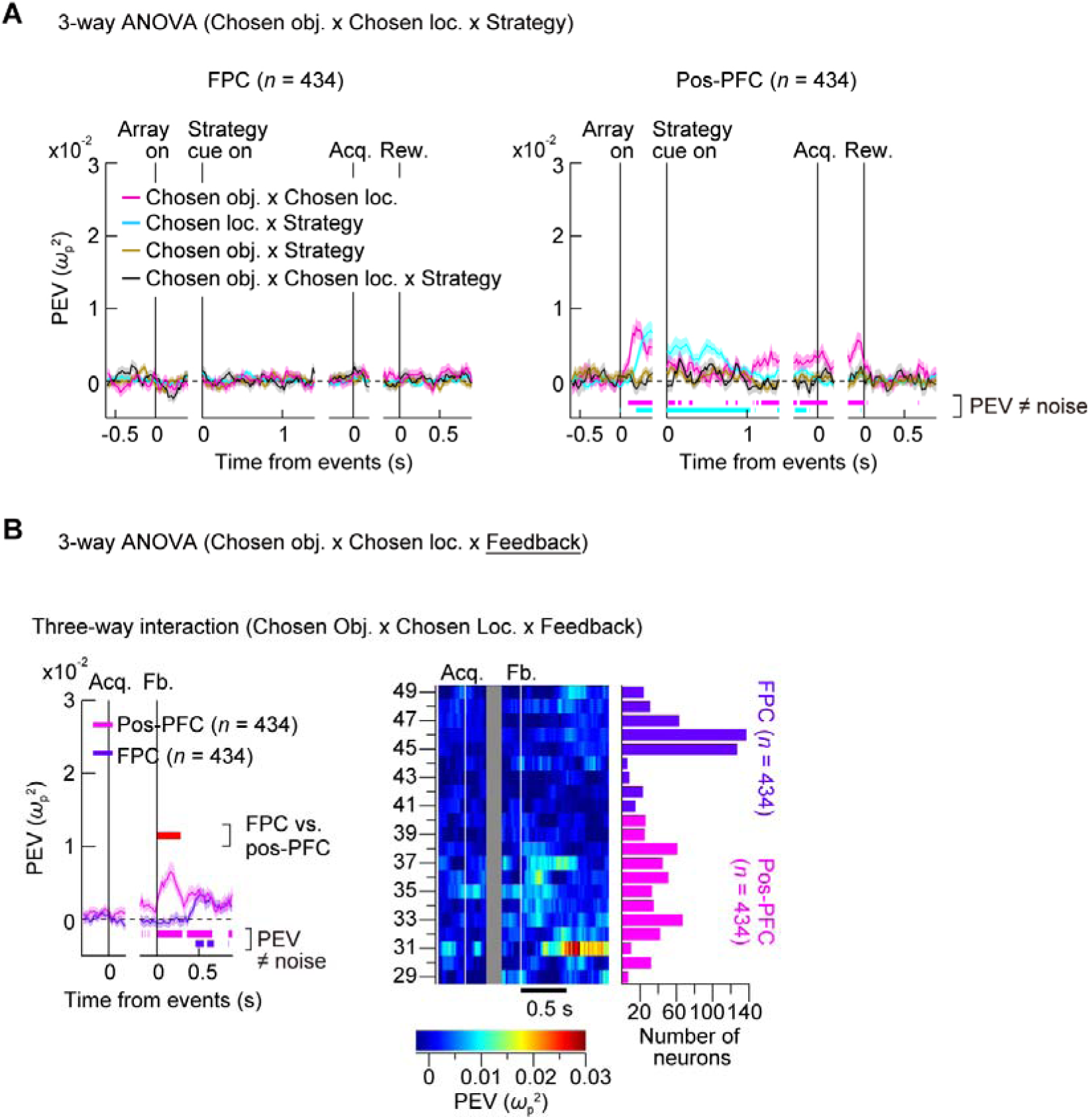
Interaction effects observed during the object CS task across the anteroposterior LPFC. (**A**) Time course of population PEV for the three first-order interaction terms and one second-order (three-way) interaction term from the 3-way ANOVA used in Figure 7H, shown separately for the FPC (left) and pos-PFC (right). Lower colored horizontal bars indicate periods of significant PEV for each term with a corresponding color. Other conventions as in Figure 7H. (**B**) Left panel: Comparison of PEV for the second-order (three-way) interaction term ([chosen object × chosen location × feedback]) between FPC (purple) and pos-PFC (magenta). Conventions as in Figure 7I. Right panel: Time course of PEV for this interaction term at each 1-mm segment along the AP axis. Conventions as in Figure 7J.

**Figure S7.**
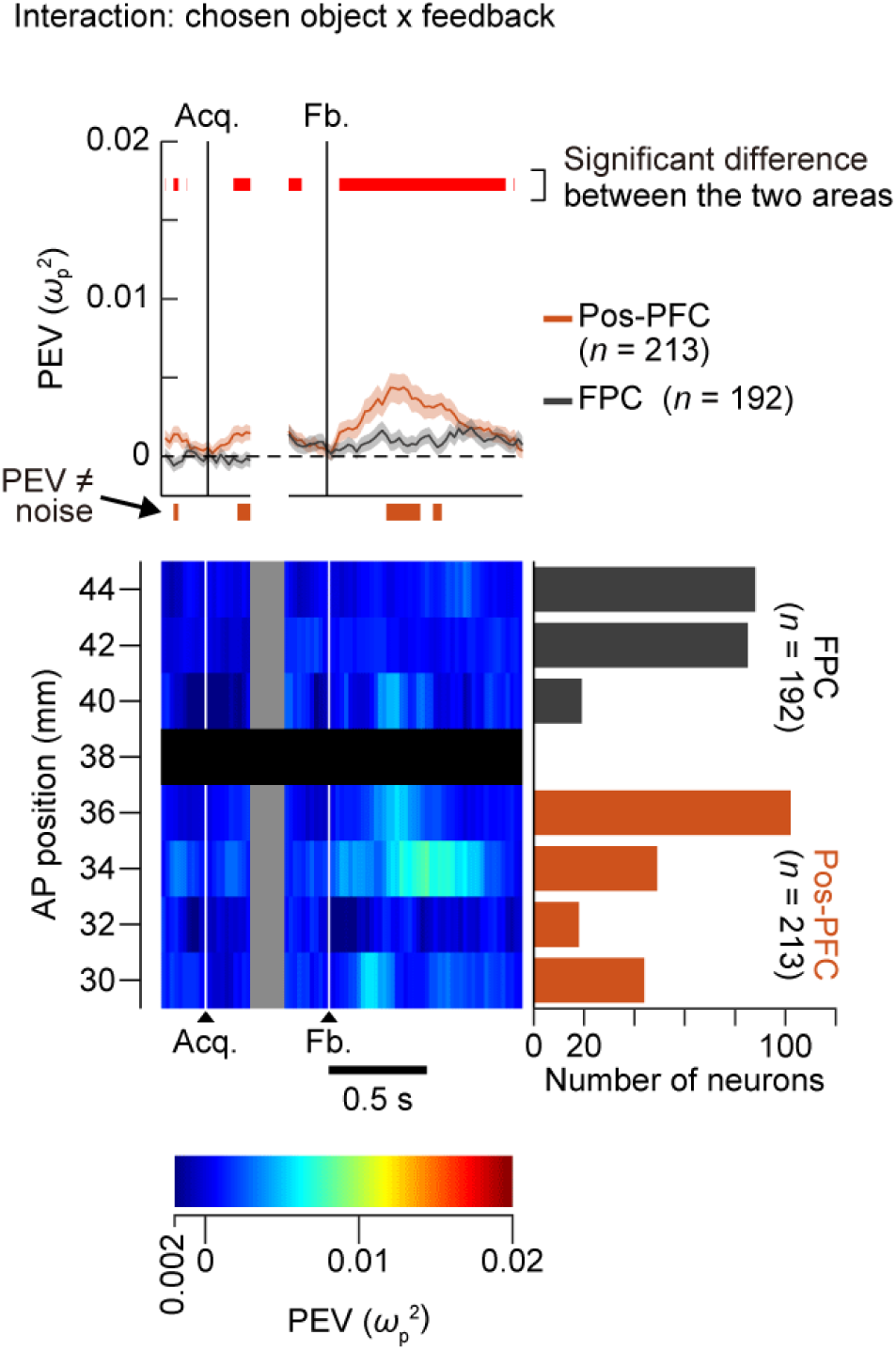
Differential encoding of decision-outcome evaluation between FPC and pos-PFC in the small-set FsL task (Experiment 4). Top panel: Time course of PEV for the critical interaction term (chosen object × feedback) in the 3-way ANOVA with factors: chosen object, feedback, and ordinal trial position. Conventions as in Figure 6D. Bottom panel: Time course of PEV for this interaction term at each 2-mm segment of recording location along the AP axis.

**Figure S8.**
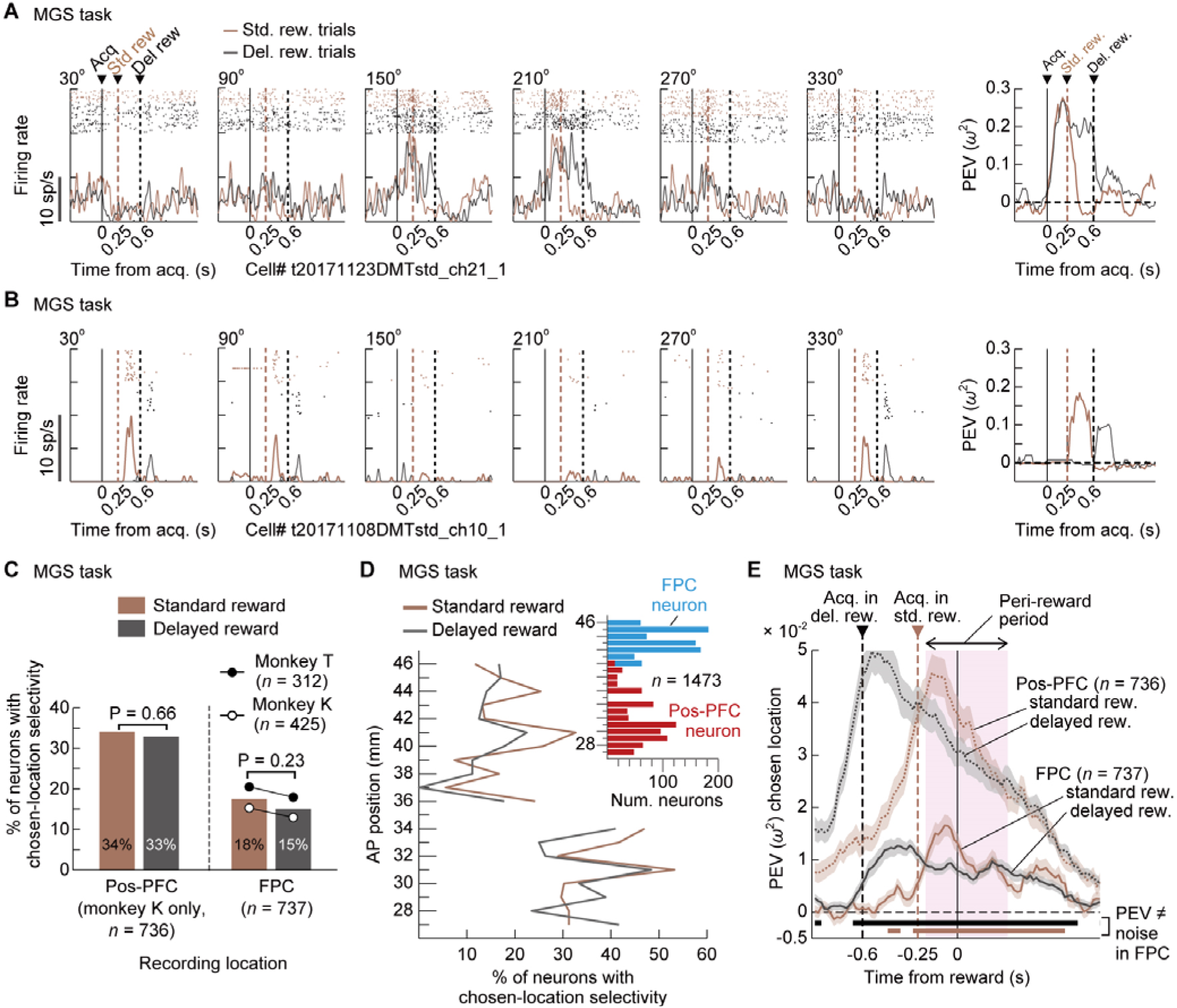
FPC neurons showed sustained chosen-location selectivity until delayed feedback even in the MGS (Experiment 1). (**A** and **B**) Raster-histograms and PEV for two representative FPC neurons in the standard (brown) and delayed reward (black) conditions in the MGS task, aligned at saccadic target acquisition (acq.). There were six possible cue locations in the MGS task (see Figure 2C). Other conventions as in Figure 8C. (**C**) Percentage of neurons with significant chosen-location (cue location) selectivity in the peri-reward period (−0.2 to 0.3 s from reward; pink shaded area in **E**) for standard (brown) and delayed reward (black) conditions in the pos-PFC (two left bars) and FPC (two right bars). Conventions as Figure 8E. (**D**) Percentage of neurons with significant chosen-location selectivity in the peri-reward period in each 1-mm segment along the AP axis. Across all 19 segments, no significant difference between the two reward conditions was observed (Fisher’s exact test, FDR-corrected). Conventions as in Figure 8F. (**E**) Time course of population-averaged PEV (mean ± s.e.m.) for chosen location in the standard (brown) and delayed reward (black) conditions, aligned at reward delivery. Results are separately shown for the FPC (solid lines) and pos-PFC (dotted lines). Lower horizontal bars indicate time periods of significant PEV in the FPC for each reward condition. Other conventions as in Figure 8G.

**Figure S9.**
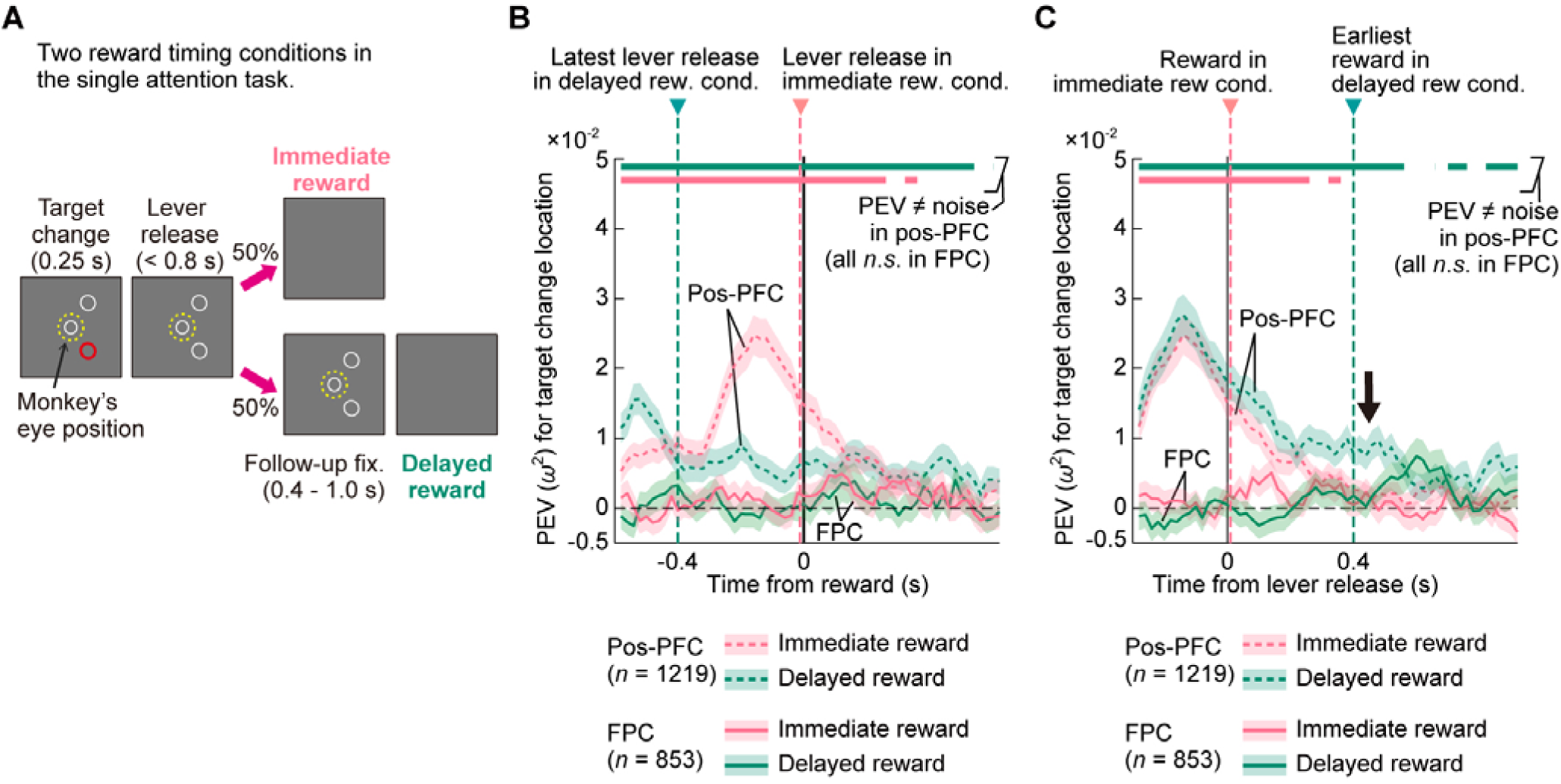
Effect of delayed feedback on neural activity in the single attention task. (**A**) Illustration of the two reward timing conditions (immediate and delayed) after correct lever release in the single attention task trials (see Figure 2A for full task description). (**B**) Time course of population-averaged PEV (mean ± s.e.m.) for target change location in pos-PFC (dashed lines) and FPC (solid lines), aligned at reward delivery. Green vertical dashed line marks the latest possible lever-release time (−0.4 s) relative to reward delivery in the delayed reward condition. Upper horizontal bars indicate periods of significant PEV in the pos-PFC (one-sample permutation test, FDR-corrected). Only pos-PFC activity showed significant PEV under both reward conditions, with no significant PEV in the FPC. (**C**) Same as in **B**, but aligned at the timing of lever release. Green vertical dashed line marks the earliest possible reward delivery time (+0.4 s) relative to lever release in the delayed reward condition. As in **B**, only pos-PFC activity showed significant PEV under both reward conditions, with no significant PEV in the FPC.

**Table S1.** Number of neurons recorded from FPC and pos-PFC by experiment and monkey.

| FPC neuron | Monkey K | Monkey T | Monkey H | Monkey Um | Total |
| --- | --- | --- | --- | --- | --- |
| Exp. 1 | 425 | 428 |  |  | 853 |
| Exp. 2 | 115 | 101 |  |  | 216 |
| Exp. 3 | 48 |  | 210 | 112 | 370 |
| Exp. 4 | 52 |  | 140 |  | 192 |
| Exp. 5 | 240 |  |  | 194 | 434 |
| Exp. 5 Suppl. <sup>a)</sup> |  |  |  | 199 | 199 |
| FPC neuron total |  |  |  |  | 2264 |
| Pos-PFC neuron | Monkey K | Monkey T | Monkey H | Monkey Um | Total |
| Exp. 1 | 736 | 483 |  |  | 1219 |
| Exp. 2 | 220 |  |  |  | 220 |
| Exp. 3 | 145 |  | 257 |  | 402 |
| Exp. 4 | 70 |  | 143 |  | 213 |
| Exp. 5 | 151 |  |  | 283 | 434 |
| Exp. 5 Suppl. <sup>a)</sup> |  |  |  | 134 | 134 |
| Pos-PFC neuron total |  |  |  |  | 2622 |

## Notes

### Competing Interest Statement

The authors have declared no competing interest.

## REFERENCES

1. P. W. Burgess, I. Dumontheil, S. J. Gilbert. The gateway hypothesis of rostral prefrontal cortex (area 10) function. Trends Cogn. Sci. 11, 290–298 (2007).

2. E. Koechlin, C. Summerfield. An information theoretical approach to prefrontal executive function. Trends Cogn. Sci. 11, 229–235 (2007).

3. D. Badre, M. D’Esposito. Is the rostro-caudal axis of the frontal lobe hierarchical? Nat. Rev. Neurosci. 10, 659–669 (2009).

4. D. E. Nee, M. D’Esposito. The hierarchical organization of the lateral prefrontal cortex. eLife 5, e12112 (2016).

5. D. Badre, D. E. Nee. Frontal cortex and the hierarchical control of behavior. Trends Cogn. Sci. 22, 170–188 (2018).

6. J. Duncan. The multiple-demand (MD) system of the primate brain: mental programs for intelligent behaviour. Trends Cogn. Sci. 14, 172–179 (2010).

7. E. A. Boschin, C. Piekema, M. J. Buckley. Essential functions of primate frontopolar cortex in cognition. Proc. Natl. Acad. Sci. U.S.A. 112, E1020–E1027 (2015).

8. F. A. Mansouri, M. J. Buckley, M. Mahboubi, K. Tanaka. Behavioral consequences of selective damage to frontal pole and posterior cingulate cortices. Proc. Natl. Acad. Sci. U.S.A. 112, E3940–E3949 (2015).

9. K. Miyamoto, R. Setsuie, T. Osada, Y. Miyashita. Reversible silencing of the frontopolar cortex selectively impairs metacognitive judgment on non-experience in primates. Neuron 97, 980–989 (2018).

10. A. Mahmoodi, C. Harbison, A. Bongioanni, A. Emberton, L. Roumazeilles, J. Sallet, N. Khalighinejad, M. F. S. Rushworth. A frontopolar-temporal circuit determines the impact of social information in macaque decision making. Neuron 112, 84–92.e6 (2024).

11. S. Tsujimoto, A. Genovesio, S. P. Wise. Evaluating self-generated decisions in frontal pole cortex of monkeys. Nat. Neurosci. 13, 120–126 (2010).

12. L. Ferrucci, S. Nougaret, F. Ceccarelli, S. Sacchetti, V. Fascianelli, D. Benozzo, A. Genovesio. Social monitoring of actions in the macaque frontopolar cortex. Prog. Neurobiol. 218, 102339 (2022).

13. S. Nougaret, L. Ferrucci, F. Ceccarelli, S. Sacchetti, D. Benozzo, V. Fascianelli, A. Genovesio. Neurons in the monkey frontopolar cortex encode learning stage and goal during a fast learning task. PLoS Biol. 22, e3002500 (2024).

14. L. Ferrucci, F. Ceccarelli, F. Londei, G. Arena, L. Elyasizad, S. Nougaret, A. Genovesio. Reward monitoring in the frontopolar cortex of macaques. Sci. Rep. 15, 1–14 (2025).

15. F. A. Mansouri, E. Koechlin, M. G. P. Rosa, M. J. Buckley. Managing competing goals—a key role for the frontopolar cortex. Nat. Rev. Neurosci. 18, 645–657 (2017).

16. K. Watanabe, S. Funahashi. Neural mechanisms of dual-task interference and cognitive capacity limitation in the prefrontal cortex. Nat. Neurosci. 17, 601–611 (2014).

17. A. Pasupathy, E. K. Miller. Different time courses of learning-related activity in the prefrontal cortex and striatum. Nature 433, 873–876 (2005).

18. R. B. Ebitz, E. Albarran, T. Moore. Exploration disrupts choice-predictive signals and alters dynamics in prefrontal cortex. Neuron 97, 450–461 (2018).

19. A. Bongioanni, D. Folloni, L. Verhagen, J. Sallet, M. C. Klein-Flügge, M. F. Rushworth. Activation and disruption of a neural mechanism for novel choice in monkeys. Nature 591, 270–274 (2021).

20. J. Achterberg, M. Kadohisa, K. Watanabe, M. Kusunoki, M. J. Buckley, J. Duncan. A one-shot shift from explore to exploit in monkey prefrontal cortex. J. Neurosci. 42, 276–287 (2022).

21. M. Ullsperger, A. G. Fischer, R. Nigbur, T. Endrass. Neural mechanisms and temporal dynamics of performance monitoring. Trends Cogn. Sci. 18, 259–267 (2014).

22. J. Grohn, U. Schüffelgen, F.-X. Neubert, A. Bongioanni, L. Verhagen, J. Sallet, N. Kolling, M. F. S. Rushworth. Multiple systems in macaques for tracking prediction errors and other types of surprise. PLoS Biol. 18, e3000899 (2020).

23. H. Tang, V. D. Costa, R. Bartolo, B. B. Averbeck. Differential coding of goals and actions in ventral and dorsal corticostriatal circuits during goal-directed behavior. Cell Rep. 38, 110189 (2022).

24. K. Watanabe, S. Funahashi. A dual-task paradigm for behavioral and neurobiological studies in nonhuman primates. J. Neurosci. Methods 246, 1–12 (2015).

25. K. Watanabe, M. Kadohisa, M. Kusunoki, M. J. Buckley, J. Duncan. Cycles of goal silencing and reactivation underlie complex problem-solving in primate frontal and parietal cortex. Nat. Commun. 14, 5054 (2023).

26. M. D’Esposito, J. A. Detre, D. C. Alsop, R. K. Shin, S. Atlas, M. Grossman. The neural basis of the central executive system of working memory. Nature 378, 279–281 (1995).

27. E. Koechlin, G. Basso, P. Pietrini, S. Panzer, J. Grafman. The role of the anterior prefrontal cortex in human cognition. Nature 399, 148–151 (1999).

28. A. Baddeley. Working memory. Science 255, 556–559 (1992).

29. M. Kadohisa, K. Watanabe, M. Kusunoki, M. J. Buckley, J. Duncan. Focused representation of successive task episodes in frontal and parietal cortex. Cereb. Cortex 30, 1779–1796 (2020).

30. W. Schultz, P. Dayan, P. R. Montague. A neural substrate of prediction and reward. Science 275, 1593–1599 (1997).

31. P. H. Rudebeck, R. C. Saunders, D. A. Lundgren, E. A. Murray. Specialized representations of value in the orbital and ventrolateral prefrontal cortex: desirability versus availability of outcomes. Neuron 95, 1208–1220 (2017).

32. S. W. Kennerley, T. E. Behrens, J. D. Wallis, Double dissociation of value computations in orbitofrontal and anterior cingulate neurons. Nat. Neurosci. 14, 1581–1589 (2011).

33. S. C. Rao, G. Rainer, E. K. Miller. Integration of what and where in the primate prefrontal cortex. Science 276, 821–824 (1997).

34. S. Tsujimoto, A. Genovesio, S. P. Wise. Neuronal activity during a cued strategy task: comparison of dorsolateral, orbital, and polar prefrontal cortex. J. Neurosci. 32, 11017–11031 (2012).

35. M. Petrides. Lateral prefrontal cortex: architectonic and functional organization. Philos. Trans. R. Soc. Lond. B Biol. Sci. 360, 781–795 (2005).

36. K. Semendeferi, E. Armstrong, A. Schleicher, K. Zilles, G. W. Van Hoesen. Prefrontal cortex in humans and apes: a comparative study of area 10. Am. J. Phys. Anthropol. 114, 224–241 (2001).

37. R. Passingham. Understanding the Prefrontal Cortex: Selective Advantage, Connectivity, and Neural Operations. (Oxford Univ. Press, 2021).

38. C. Amiez, J. Sallet, C. Giacometti, C. Verstraete, C. Gandaux, V. Morel-Latour, A. Meguerditchian, F. Hadj-Bouziane, S. Ben Hamed, W. D. Hopkins, E. Procyk, C. R. E. Wilson, M. Petrides. A revised perspective on the evolution of the lateral frontal cortex in primates. Sci. Adv. 9, eadf9445 (2023).

39. J. Sallet, R. B. Mars, M. P. Noonan, F.-X. Neubert, S. Jbabdi, J. X. O’Reilly, N. Filippini, A. G. Thomas, M. F. Rushworth. The organization of dorsal frontal cortex in humans and macaques. J. Neurosci. 33, 12255–12274 (2013).

40. S. T. Carmichael, J. L. Price. Connectional networks within the orbital and medial prefrontal cortex of macaque monkeys. J. Comp. Neurol. 371, 179–207 (1996).

41. A. Goulas, H. B. Uylings, P. Stiers. Mapping the hierarchical layout of the structural network of the macaque prefrontal cortex. Cereb. Cortex 24, 1178–1194 (2014).

42. T. Kojima, H. Onoe, K. Hikosaka, K. I. Tsutsui, H. Tsukada, M. Watanabe. Default mode of brain activity demonstrated by positron emission tomography imaging in awake monkeys: higher rest-related than working memory-related activity in medial cortical areas. J. Neurosci. 29, 14463–14471 (2009).

43. G. Paxinos, M. Petrides, H. C. Evrard. The Rhesus Monkey Brain in Stereotaxic Coordinates. (Academic Press, 2023).

44. P. H. Rudebeck, T. E. Behrens, S. W. Kennerley, M. G. Baxter, M. J. Buckley, M. E. Walton, M. F. Rushworth. Frontal cortex subregions play distinct roles in choices between actions and stimuli. J. Neurosci. 28, 13775–13785 (2008).

45. E. Koechlin. Frontal pole function: what is specifically human? Trends Cogn. Sci. 15, 241 (2011).

46. N. P. Friedman, T. W. Robbins. The role of prefrontal cortex in cognitive control and executive function. Neuropsychopharmacology 47, 72–89 (2022).

47. E. D. Boorman, T. E. Behrens, M. W. Woolrich, M. F. Rushworth. How green is the grass on the other side? Frontopolar cortex and the evidence in favor of alternative courses of action. Neuron 62, 733–743 (2009).

48. J. P. Stroud, K. Watanabe, T. Suzuki, M. G. Stokes, M. Lengyel. Optimal information loading into working memory explains dynamic coding in the prefrontal cortex. Proc. Natl. Acad. Sci. U.S.A. 120, e2307991120 (2023).

49. C. B. Drucker, M. L. Carlson, K. Toda, N. K. DeWind, M. L. Platt. Non-invasive primate head restraint using thermoplastic masks. J. Neurosci. Methods 253, 90–100 (2015).

50. R. Tanaka, K. Watanabe, T. Suzuki, K. Nakamura, M. Yasuda, H. Ban, K.-i. Okada, S. Kitazawa. An easy-to-implement, non-invasive head restraint method for monkey fMRI. NeuroImage 285, 120479 (2024).

51. A. Kuznetsova, H. Rom, N. Alldrin, J. Uijlings, I. Krasin, J. Pont-Tuset, S. Kamali, S. Popov, M. Malloci, A. Kolesnikov, T. Duerig, V. Ferrari. The open images dataset v4: Unified image classification, object detection, and visual relationship detection at scale. Int. J. Comput. Vis. 128, 1956–1981 (2020).

52. A. Fedorov, R. Beichel, J. Kalpathy-Cramer, J. Finet, J. C. Fillion-Robin, S. Pujol, R. Kikinis. 3D Slicer as an image computing platform for the Quantitative Imaging Network. Magn. Reson. Imaging 30, 1323–1341 (2012).

53. M. R. Riley, X. L. Qi, X. Zhou, C. Constantinidis. Anterior-posterior gradient of plasticity in primate prefrontal cortex. Nat. Commun. 9, 3790 (2018).

54. Y. Benjamini, Y. Hochberg. Controlling the false discovery rate: A practical and powerful approach to multiple testing. J. R. Stat. Soc. Ser. B 57, 289–300 (1995).

55. D. Cousineau. Confidence intervals in within-subject designs: A simpler solution to Loftus and Masson’s method. Tutor. Quant. Methods Psychol. 1, 42–45 (2005).

56. R. D. Morey. Confidence intervals from normalized data: A correction to Cousineau (2005). Tutor. Quant. Methods Psychol. 4, 61–64 (2008).

